# Simple and complex interactions between sleep-wake driven and circadian processes shape daily genome regulatory dynamics in the mouse

**DOI:** 10.1101/677807

**Authors:** Charlotte N. Hor, Jake Yeung, Maxime Jan, Yann Emmenegger, Jeffrey Hubbard, Ioannis Xenarios, Felix Naef, Paul Franken

**Affiliations:** Centre for Integrative Genomics, University of Lausanne, Batiment Genopode, Quartier Sorge, CH-1015 Lausanne, Switzerland; Institute of Bioengineering, School of Life Sciences, Ecole Polytechnique Fédérale de Lausanne, EPFL SV IBI UPNAE, AAB 040, Station 15, CH-1015 Lausanne, Switzerland; Swiss Institute of Bioinformatics, Batiment Genopode, Quartier Sorge, CH-1015 Lausanne, Switzerland

**Keywords:** circadian, sleep, gene expression, epigenetics, long-term effects, exposure

## Abstract

The timing and duration of sleep results from the interaction between a sleep-wake driven, or homeostatic, process (S) and a circadian process (C), and involves changes in gene expression and genomic regulation. Unraveling the respective contributions of S and C, and their interaction, to transcriptional and epigenomic regulatory dynamics requires sampling over time under unperturbed conditions and conditions of perturbed sleep. Here, we profiled mRNA expression and chromatin accessibility in the cerebral cortex of mice over a three-day period, including a 6-hour sleep deprivation (SD) on day two. Mathematical modeling established that a large proportion of rhythmic genes are actually governed by Process S with varying degrees of interaction with Process C, sometimes working in opposition. Remarkably, SD causes long-term effects on gene expression dynamics, outlasting phenotypic recovery, most strikingly illustrated by a dampening of the oscillation of most core clock genes, including *Bmal1*, suggesting that enforced wakefulness directly impacts the molecular clock machinery. Chromatin accessibility proved highly plastic and dynamically affected by SD. Distal regions, rather than promoters, display dynamics corresponding to gene transcription, implying that changes in mRNA expression result from constantly accessible promoters under the influence of distal enhancers or repressors. *Srf* was predicted as a transcriptional regulator driving immediate response, suggesting that *Srf* activity mirrors the build-up and release of sleep pressure. Our results demonstrate that a single, short SD has long-term aftereffects at the genomic regulatory level. Such effects might accumulate with repeated sleep restrictions, thereby contributing to their adverse health effects.

**Significance statement:** When and how long we sleep is determined by the time-of-day and how long we have been awake, which are tracked molecularly by a circadian and a sleep-wake driven process, respectively. We measured the long-term consequences of a short-term sleep deprivation (SD) on gene expression and regulation in the mouse brain, and used mathematical models to determine the relative contributions of the circadian and sleep-wake driven processes. We find that many genes, including most of the genes that constitute the molecular circadian clock, are perturbed by SD long after the mice ceased showing behavioral signs of sleep loss. Our results have implications for human health, given the high prevalence of insufficient and poor quality sleep in our contemporary society.

## Introduction

According to the two-process model (1, 2), sleep regulation results from an interaction between the sleep homeostatic process (Process S) and the circadian process (Process C). The sleep homeostat keeps track of pressure for sleep as it increases during wake and decreases during sleep, while the circadian process dictates the optimal time-of-day for sleep to occur. Their fine-tuned interaction assures optimal timing, duration and quality of both wakefulness and sleep, and even minor changes in either of these processes or their alignment cause performance decrements and clinically significant sleep disruption (3, 4).

The circadian clock is described as self-sustained 24h oscillations involved in a variety of physiological processes and behaviors such as sleep (3, 5). It is encoded molecularly through negative feedback loops involving the core clock genes, which are capable of generating oscillations in constant environmental conditions, *i.e.* in the absence of periodically occurring time cues such as the light-dark cycle (6). However, this apparent autonomy does not inevitably imply that the expression of all genes displaying a rhythm with a period of 24-hours is directly driven by the circadian clock. For example the light-dark cycle, besides entraining the circadian clock, directly influences many physiological and behavioral processes (7). Also, the rhythmic organization of sleep-wake behavior and associated feeding and locomotion directly drives gene expression (8). Disentangling the respective contributions of the circadian and sleep-wake driven processes is experimentally challenging and has been addressed by methods suppressing one component (*e.g.* surgical or genetic ablation of circadian oscillators) or uncoupling their relationship through forced desynchrony or sleep deprivation (SD) (3, 9).

SD experiments aiming at identifying genes associated with the sleep homeostatic process follow the rationale that causing mice to stay awake during a time when they normally sleep will induce an acute response in sleep-wake driven genes. Indeed, studies comparing gene expression levels immediately after SD with controls collected at the same time-of-day have identified many differentially expressed genes (10–13), and a few studies have probed the punctual effect of SD at different times of the 24h cycle in mice (13, 14), or expression dynamics in blood during SD in humans (15, 16). However, assessing the respective contributions of the two processes requires measuring gene expression over multiple time points, not only under SD, *i.e.* enforced waking, but also under spontaneous sleep-wake dynamics pre- and post-SD. Furthermore, to systematically link temporal gene expression to the sleep-wake distribution and/or circadian clock, the analysis should consider the entire time series, rather than only pair-wise differential comparisons. Finally, the regulatory mechanisms underlying such dynamics are largely unexplored (17), particularly in this kind of dynamic context.

To systematically investigate the gene expression dynamics caused by one acute SD episode, as well as the underlying regulatory events, we measured chromatin accessibility alongside mRNA expression in the cerebral cortex of adult C57BL6/J mice over 24 hours before, during, and over 48 hours following one 6-hour session of total SD, as well as 7 days after the intervention. We modeled the entire time series based on the assumptions of the two-process model to objectively assess whether the mRNA accumulation dynamics of each cortically expressed gene follow Process S, Process C or a combination thereof. This setting allowed us to characterize the temporal dynamics of the consequences of SD on gene expression and regulation, and dissect the interaction between Processes S and C. Moreover, we identified genomic regulatory elements implicated in the transcriptional response to sleep loss by exploring the hitherto understudied epigenetic landscape of sleep (18).

## Results

### Behavioral response and recovery after sleep deprivation

We observed the typical distribution of sleep over 24 hours in baseline, with mice spending most of the light period asleep, while being predominantly awake during the dark period (Fig 1B, bottom). Delta power in NREM sleep (Fig 1B, top), an EEG-derived variable considered to reflect sleep ‘pressure’ (Process S), was high after spontaneous waking in the dark phase and decreased during the light phase, and we observed the well-known effects of SD, namely an increase in delta power in the 45 minutes immediately following SD and a rebound of time spent in NREM sleep observed during the first 12h of recovery (T30-T42). We found that values for NREM sleep no longer significantly deviated from baseline levels already after T42 (Fig 1B bottom, black line). During the dark phase (T36-48), delta power even dropped below the levels reached at this time during baseline, likely as a consequence of the increased time spent in NREM sleep during the first 12h after SD. REM sleep was affected in the same manner as NREM sleep (Fig 1B, bottom, green line).

**Fig 1.**
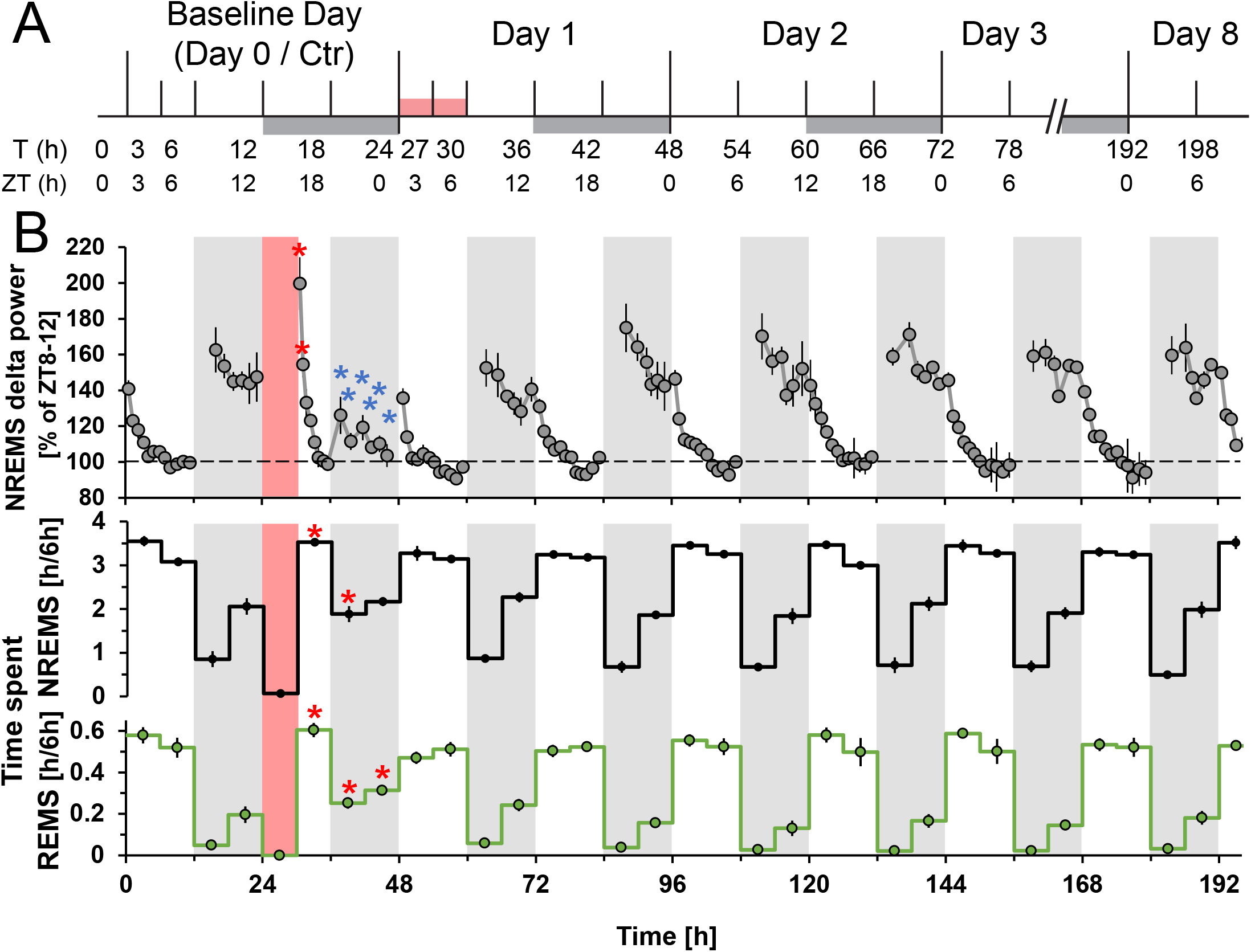
Study design and long-term effects of SD on sleep behavior and EEG delta power. **(A)** Tissue collection schedule with time in hours from beginning of the experiment (T) and corresponding zeitgeber time (ZT). White and grey bars: 12h:12h light/dark cycle. Red bar: SD. **(B)** Long-term effects of SD on NREM sleep delta power (1-4Hz, top), and NREM and REM sleep quantity (bottom, black, respectively green lines). Mean delta power values (± SEM) are expressed as the percentage of intra-individual deviations from the time interval in baseline with the lowest overall power (ZT8-12, average across 2 days). Asterisks denote significant increases (red) and decreases (blue) compared to baseline (t-test, p<0.05, n=6). White and grey shading: 12h:12h light/dark cycle. Red shaded area: SD.

We asked whether the fast reversal of the phenotype in the EEG data would be paralleled by changes at the gene expression and regulatory levels in the cerebral cortex, or whether novel molecular dynamic patterns could be observed. We therefore measured and analyzed the temporal dynamics of transcriptomes and chromatin accessibility over a total of 78 hours, including baseline, SD and recovery.

### Sleep-wake history is the main driver of transcriptome dynamics

We first examined the detected fraction of the transcriptome (13’842 genes) using principal component analysis (PCA, Fig 2A). We observed that samples formed three groups along the first principal component (PC1) axis. The left-most group gathered time points during the light phase of the LD cycle where mice generally spend more time asleep, while the middle group represented time points during the dark phase when mice are predominantly awake (*i.e.* the former group spent more time asleep prior to sampling than the latter). This separation could evoke that PC1 separates samples according to time of day, however this notion is challenged by the shift towards the right of the samples taken at ZT3 and ZT6 during SD (T27 and T30), *i.e.* in the complete absence of sleep, suggesting that the PC1 axis follows (from left to right) increased time spent awake prior to sampling rather than zeitgeber time (ZT).

**Fig 2.**
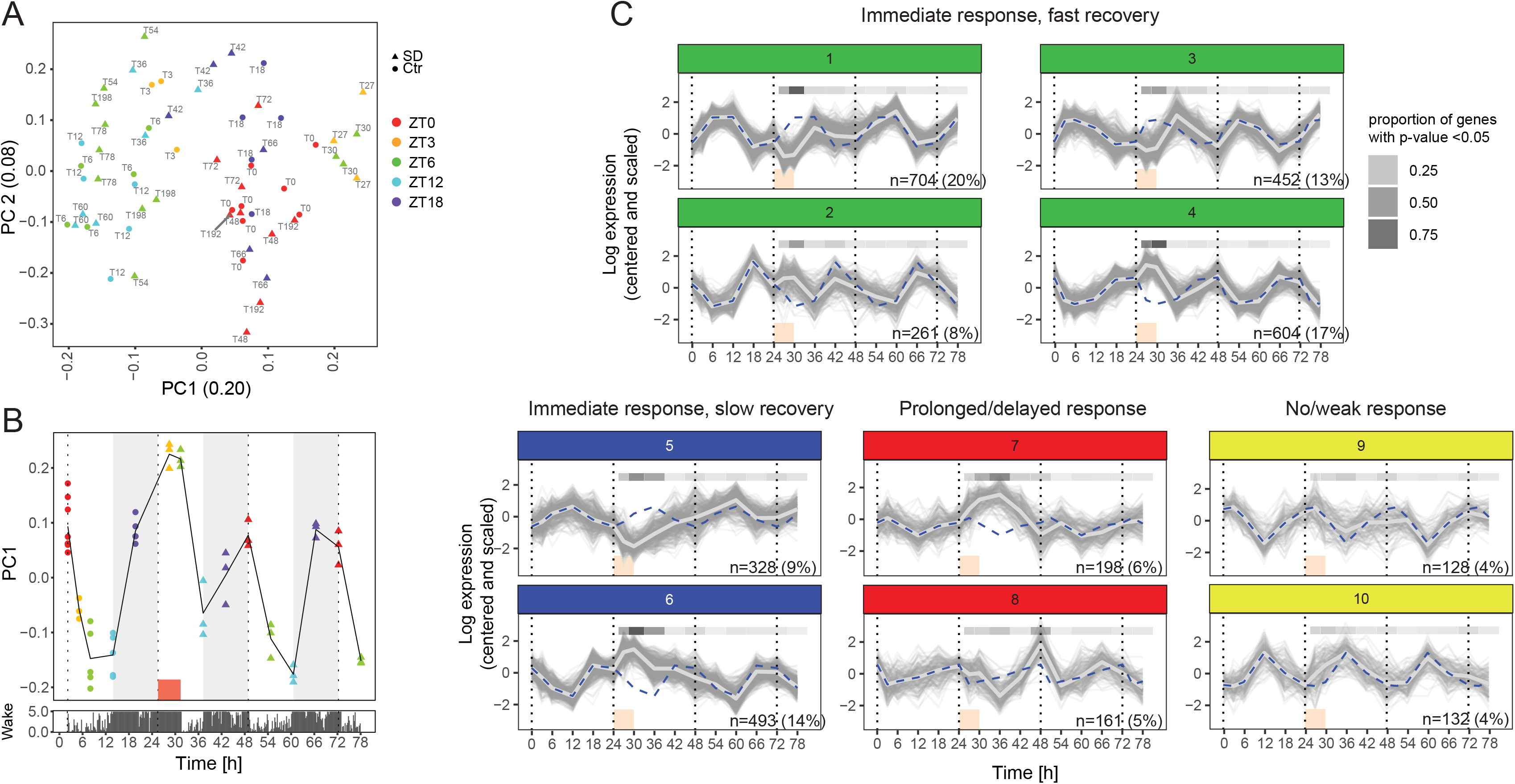
Sleep-wake history is the main driver of transcriptome dynamics. **(A)** Principal component analysis of the expression of the 13’842 detected genes. The number in parenthesis in the axis label denotes the fraction of the variance explained by the component. Colors denote zeitgeber time (ZT0-ZT12: light period, ZT12-ZT0: dark period). Baseline samples are represented by discs and samples collected during or after SD by triangles. Text labels denote time of sample collection according to the experimental design (see Fig 1A). **(B)** First principal component plotted over time (top). Time spent awake over time averaged across 12 mice (bottom), with y-axis denoting the number of minutes the mice spent awake over the last 5 minutes. **C.** K-means clusters of the 3461 genes with statistically significant temporal gene expression (FDR adjusted *p*-value < 0.001, likelihood ratio test). Blue dashed line: average of the cluster under baseline, repeated for comparison over the three days of the experiment. Light grey thick line: cluster average. Red box: SD. Grey shaded bar at the top of each graph: proportion of genes with *p*-value <0.05 according to a likelihood ratio test between SD and baseline conditions at the same ZT.

To illustrate the sleep-wake-driven dynamics underlying PC1, we overlaid PC1 with the average amount of waking over time (Fig 2B). PC1 increased during periods of waking, decreased during periods of sleep, and, importantly, reached its maximum during SD, in a pattern strongly reminiscent of Process S and EEG delta power ((19) and Fig 1B, top). PC1 thus reflects the amount of sleep prior to sample collection and highlights the pervasive impact of sleep-wake distribution on gene expression, which we further explore below.

### Clustering of mRNA temporal profiles highlights diverse response and recovery kinetics

To uncover and classify general temporal patterns in our data, we first performed an exploratory analysis using k-means clustering. With this unsupervised clustering, we grouped the temporal expression of 3461 genes displaying statistically robust temporal variation from T0 to T78 (FDR-adjusted *p*-value < 0.001, see Methods). This analysis on a conservatively selected subset of genes aimed to uncover broad dynamics in the dataset. We observed distinct profiles of response to SD and subsequent recovery by comparing the cluster average of T24-T78 with the baseline day (T0-T18, Fig 2C, light grey line and blue dashed line, respectively). Here, “response” refers to temporal patterns from T24 to T78 that deviate from baseline at corresponding ZT times (e.g., T30 and T54 are compared with T6; T36 and T60 are compared with T12, etc.). “Recovery” refers to patterns that return towards baseline levels. Genes in clusters 1-6 displayed an immediate response, with many showing marked differences detectable already during and at the end of SD, *i.e.* at T27 and T30 (Fig 2C). In cluster 7, the response progresses until T36, after which reversal takes place. This “prolonged” response is to differentiate from a “delayed” response as in cluster 8, where the first significant difference to baseline is visible 6 hours after the end of SD.

At the level of recovery, clusters 1-4 show a fast recovery, where baseline levels are reached already at T36, making them reminiscent of delta power dynamics. Meanwhile, clusters 5 and 6 contain genes that revert more slowly, reaching baseline at T42 in a pattern paralleling the recovery dynamics of time spent asleep. Cluster 7 also displays slow recovery, with baseline levels attained 12 hours after the peak response at T36. In cluster 8 we observe a distinctive recovery pattern in the form of an increase at T48 following the initial downregulation at T36. Finally, clusters 9 and 10 showed a prominent 24-hour rhythm with subtle, if any, perturbation by SD (mean *p*-values across genes > 0.24).

Generally, the fast response and fast recovery dynamics, together with a direction of change opposite to what is expected by time-of-day, suggests that these genes are sleep-wake driven, *i.e.* genes that usually go up when the mouse is predominantly asleep are downregulated during SD (clusters 2, 4, 6) and *vice versa* (clusters 1, 3, 5). Also, the slower dynamics of response and recovery (cluster 7, 8) suggests that the effects of SD can also occur downstream of the immediate response and last beyond the exposure itself, and suggests that the time required for molecular recovery exceeds the time for phenotypic recovery.

### Modeling temporal transcriptome dynamics

To better characterize and distinguish these expression dynamics, we sought, given our observations, to model gene expression over the entire time course including baseline, SD and the aftermath of SD, taking into account that the response and recovery can be sleep-wake driven, circadian or a mixture of both. Also, because rhythmicity can be suppressed or altered during mistimed or restricted sleep (13, 15, 16, 20), we included the possibility that it could remain perturbed after the end of the SD.

Explicitly modeling the temporal dynamics of mRNA profiles can offer advantages over unsupervised methods such as the k-means clustering implemented above. Indeed, the parameters of a model can give biological insights into the underlying dynamics, competing models can be systematically compared, and explicit hypotheses can be tested. Thus, modeling will unify dynamics that appear in separate clusters (e.g., Cluster 1 and 4 may both have sleep-wake driven genes), and differentiate dynamics that appear in the same cluster (e.g., Cluster 9 contains both SD-resistant and SD-sensitive dynamics, see Fig S1A).

We thus devised 6 models to explain the gene expression dynamics of the full transcriptome (n=13’842 detected genes) (Fig S1B): (1) constant or ‘flat’ model (F); (2) sleep-wake history modeled from sleep-wake data (S, in analogy to Process S in the 2-process model (1)); (3) cosine dynamics with a 24-hour rhythm (C); (4) cosine with amplitude change after SD (C_A_); (5) sleep-wake + cosine (S+C); (6) sleep-wake + cosine with amplitude change (S+C_A_). To select the best among competing models, we used the Bayesian Information Criterion (BIC) to balance model fit and model complexity, which we transformed into model weights *w* (see Methods). For each gene, the sum of the weights for all 6 models equals 1, and the model with the highest weight is assigned to the gene. Each gene is assigned one of the 6 models. In the example of the core clock gene *Nr1d1* (*RevErb*α, Fig 3A), the selected model was cosine with amplitude change after SD (model C_A_, represented as a bold line), due to a very high weight *w*=0.977. Indeed, the baseline pattern (from T0 to T24) is consistent with a circadian oscillation, the amplitude of which is significantly reduced after SD and, surprisingly, not re-established by T78. All fits are presented in Table S4.

**Fig 3.**
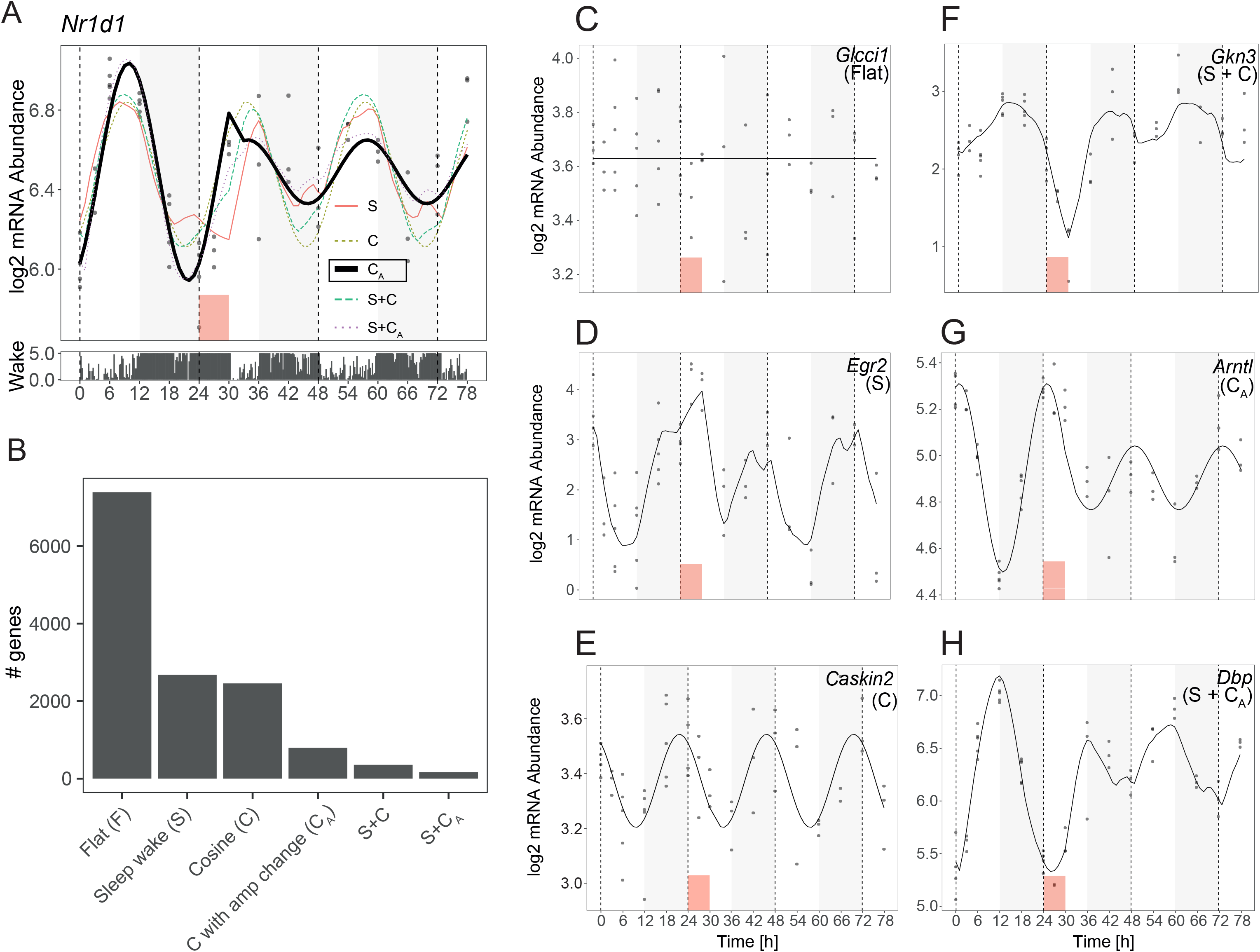
Modeling temporal transcriptome dynamics. **(A)** Example of model fitting on *Nr1d1*. Dots: RNA level data points. Bold line: best fitting model (here C_A_ with *w*=0.977). Temporal EEG data as in Fig 2. Red box: SD. **(B)** Number of genes per model. (**C-H**) Examples of a gene fit to each model.

### Model S recapitulates known sleep-wake driven genes and closely parallels EEG delta power dynamics

We summarized the genes assigned to each model genome-wide (Fig 3B), and found that, out of all temporal models (*i.e.* excluding the flat model which fit 7391 genes, example Fig 3C), the sleep-wake driven model (model S) had the largest number of genes assigned to it (2677 genes, example Fig 3D), consistent with the interpretation of PC1 reflecting sleep-wake history and the predominance of fast response dynamics in our cluster analysis (clusters 1-6). Analyzing the parameters of genes associated with model S, we found that the fitted time constants describing model S corresponding to wake (*τ*_*w*_, median = 7.05 h) and sleep (*τ*_*s*_, median = 1.68 h) were strikingly close to the dynamics of EEG delta power found in (19) (8.0h, respectively 1.8h) for this inbred strain. These dynamics closely resemble an immediate early gene (IEG) response, raising the possibility that EEG delta power is linked to, or even preceded by, a molecular change in the brain.

Model S included genes previously described as affected at the end of SD such as *Egr2*, *Arc*, *Fos* and *Cirbp* (12, 13, 21) (Fig 3D and Fig S2 pages 1-3, all *w*>0.833), but also, surprisingly, the core clock genes *Clock* and *Npas2* (*w*=0.787, resp. *w*=0.527, Fig S2 pages 4-5). Generally, we found the overwhelming majority of known sleep-wake driven genes to be correctly assigned by our model selection method. For example, the dynamics of 72 out of 75 (96%) genes previously described as sleep-wake driven (21) were affected by SD and fully or partially explained by the sleep-wake data (*i.e.* assigned to models S, S+C and S+C_A_, Fig 4A, *p*-value = 1.2e-14, chi-squared test). Similarly, we found that 181/207 (87%) genes previously described as affected by SD at any time of day in whole brain (Table S5 in Ref. (13)) were also inferred to be affected by SD (*i.e.* assigned to the same models) in our model selection (Fig 4B, *p*-value = 2.6e-21, chi-squared test). Consistently, genes that were upregulated during waking all had their maximum expression during the baseline dark period, while those downregulated during waking peaked during the baseline light period (Fig 5A).

**Fig 4.**
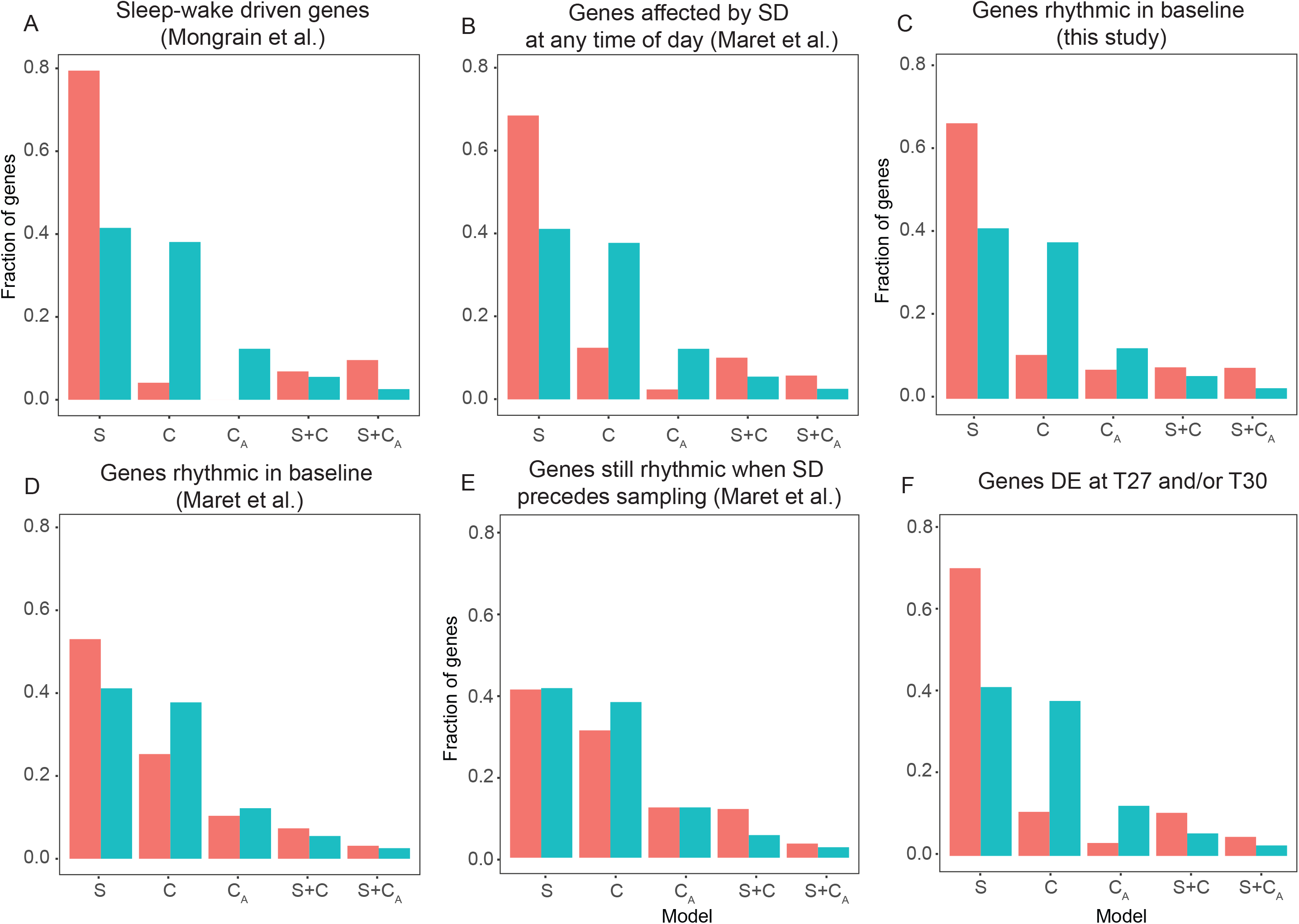
Repartition of gene sets in the different models, contrasted with all transcripts detected. Red bars: target set. Blue bars: all detected genes. Enrichment statistic by chi-square test. **(A)** 75 sleep-wake driven genes from (21), *p*-value = 1.2e-14. **(B)** 207 genes affected by SD at any time of day (Table S5 in (13)), *p*-value = 2.6e-21. **(C)** 862 rhythmic genes during baseline Day 0 (this study, *p*-value = 2.4e-76). **(D)** 918 genes rhythmic under baseline conditions (Table S3 in (13)), *p*-value = 2.3e-14. **(E)** 260 genes still rhythmic when sample collection was preceded by 6h SD (Table S4 in (13)), *p*-value = 0.00024. **(F)** 2863 genes differentially expressed at T27 and/or T30 (*i.e.* the union of T27 vs. T3 and T30 vs. T6, *p*-value = 1.2e-237).

**Fig 5.**
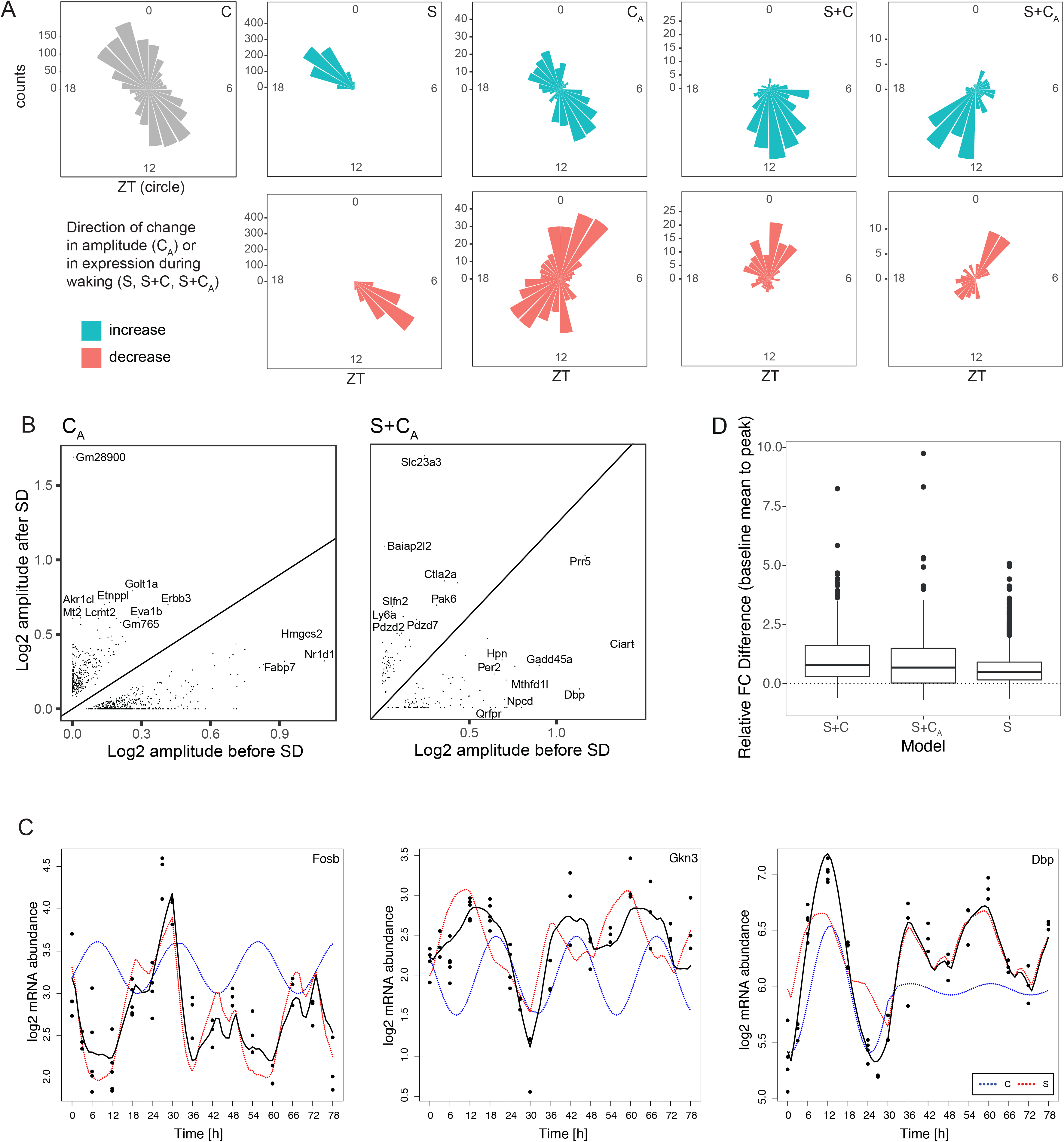
Contribution of the S and C components to phase and amplitude of oscillating genes under baseline and after SD. **(A)** Phase maps of non-flat model genes. Each cone reflects the number of genes peaking during one hour around the ZT clock. Non-C model genes are divided according to the direction of change of the amplitude (C_A_), or the direction of the change in expression happening under wakefulness (S, S+C, S+C_A_), *i.e.* the sign of the difference between the expression value given by the model after long wakefulness episodes minus that after long sleep episodes. **(B)** Scatter plots of the log2 amplitude before and after SD for models C_A_ and S+C_A_. **(C)** Contribution of the S and C components (red and blue dotted curves, respectively) to the overall temporal gene expression profile (solid black curve) of three example genes. **(D)** Difference between the highest point before and after SD, both compared to the baseline average, for models S, S+C and S+C_A_. **(E)** Number of differentially expressed genes at each time point.

### mRNA expression of all but one core clock gene is sensitive to SD

Following closely, the cosine model (model C) was the third most abundant category with 2457 genes. This model gathers genes the oscillation of which is largely unaffected by SD as illustrated by the top fit *Caskin2* (*w*=0.880, Fig 3E). The genes with the largest amplitude showing 24-hour oscillations in gene expression resistant to SD were *Sgk1*, a glucocorticoid regulated kinase, and *Cldn5*, a principal tight junction protein in the blood-brain barrier (Fig S2 pages 6-7), reaching their peak just before the light-to-dark and dark-to-light transitions, respectively. Generally, examining the oscillation patterns of this group of genes, we find that the phases (*i.e.* time at which expression peaks) of model C genes are not random, but tend to accumulate in the second half of the light or dark phases (ZT10 and ZT22, Fig 5A).

Interestingly, our algorithm assigned *Cry1* as the only core clock gene in this model, with model C_A_ second in explaining its mRNA dynamics (*w*=0.661 and *w*=0.336 respectively, Fig S2 page 8). The other clock genes, instead of fitting model C as might be expected, were assigned to model S (see above), the altered amplitude model C_A_ or the more complex models (models S+C and S+C_A_, see below). Clock genes assigned to model C_A_ (which contained in total 794 genes) had dampened amplitudes after SD, and encompassed the example *Nr1d1/Rev-erbα* (Fig 3A), as well as *Arntl* (*Bmal1*, Fig 3G), *Per3*, *Cry2* and *Nr1d2* (*RevErbβ*, Fig S2 pages 9-11). Except for *Cry2*, where the top weights were *w*(C_A_)=0.645 and *w*(C)=0.347, the model C weight *w*(C) of the other clock genes was negligible (highest *w*(C)=0.0003 for *Nr1d1*), meaning the clock genes were assigned to C_A_ either unequivocally, or with a close call to the more complex models S+C and S+C_A_ (Table S4). Model C_A_ also contained genes with increased amplitudes (331/794; 42%) after SD, such as *Erbb3*, *Eva1b*, *Zfp473* and *Akr1cl* (Fig 5B, Fig S2 pages 12-15). Interestingly, the phases of expression differed between the genes with increased vs. decreased amplitudes, the former group having a similar distribution of phases to those of model C genes (Fig 5A).

### Oscillating transcripts in baseline are often sleep-wake driven

We next asked if genes with rhythmic expression in our mouse cortex dataset were affected by SD. We therefore applied harmonic regression to the baseline Day 0 time points, which yielded a set of 862 oscillating genes (FDR-adjusted *p*-value < 0.05). Strikingly, we found that the majority of these genes (578, 67%) were assigned to the S model, and that the C model was impoverished in these genes (Fig 4C, *p*-value = 2.4e-76, chi-squared test). Again, we compared these results to a published set of 918 genes (corresponding to 2032 probesets) rhythmic under baseline conditions in the whole brain (Table S3 in (13)) and found a similar repartition (Fig 4D, *p*-value = 2.3e-14, chi-squared test), which suggests that many of the genes that appear rhythmic in undisturbed conditions oscillate as a consequence of sleep and wake (which itself is rhythmic) rather than because of the circadian clock directly. We also examined the repartition of another set from the same study, namely 391 probesets which were still rhythmic when sampling was preceded by 6 hours SD (Table S4 in (13)). We expected these genes to be resistant to SD and classify mainly in model C, however, we found that their repartition among models was only marginally different from our complete set, with even a slight impoverishment of model C and enrichment of model S+C (Fig 4E). This indicates that the rhythmicity observed in that study was due to other factors besides circadian, such as the presence of a light-dark cycle, differences in SD side-effects (e.g. stress) when performed at different times of day, or differences in conditions prior to SD (such as time-spent-awake).

### Sleep and the circadian process can work in opposition to limit oscillation amplitudes in baseline

Model S+C (357 genes) incorporated the sleep-wake history and time-of-day to output a strong response to SD while maintaining modest fold changes during baseline (*e.g. Gkn3*, *Per1* and *Fosb*, Fig 3F and Fig S2 pages 16-17). Indeed, this model, together with its altered-amplitude counterpart (S+C_A_, see below) allowed to explain complex temporal patterns, notably due to interactions between the S and C models, additive in the log scale. For genes upregulated during SD, both the S and C components increased concurrently during SD, but discordantly in baseline (e.g. *Fosb*, Fig 5C). Similarly, for genes downregulated during SD, the S and C components decreased concurrently during SD, but discordantly in baseline (e.g. *Gkn3*, Fig 5C). The discordant action caused dynamics in mRNA levels to be limited during baseline, while the concordant action during SD allows large fold changes relative to baseline. In comparison, in model S where the buffering by the C component is absent, the difference in fold change from the highest point after SD and the highest point in baseline, both compared to the baseline average, was smaller than for models S+C and S+C_A_ (Fig 5D), meaning that expression changes in model S genes are similar whether wakefulness is spontaneous or enforced, and regardless of time of day. We note that S+C genes tend to peak around the light-to-dark and dark-to-light transitions, a shift in comparison with model C and model S genes (Fig 5A).

The observation of this S+C interaction provides an intriguing parallel with human and primate studies of cognitive performance under forced desynchrony or SD protocols, where it was found that the phase of the circadian wake-promoting signal is timed in such a way that it opposes the sleep-wake dependent accumulation of sleep propensity and peaks in the hour prior to habitual sleep onset. This timing is essential for maintaining high and stable levels of attention and cognitive performance during the day as well a consolidated period of sleep during the night (22–27).

### SD represses the C component of genes with complex dynamics, leaving them predominantly under the control of Process S

The most complex of our models, model S+C_A_ (166 genes, example *Dbp*, Fig 3H), incorporated sleep-wake history, time of day and altered amplitudes to model the response to SD and subsequent change in amplitudes after SD. In this model, we see that the additive dynamic process observed in model S+C (see above) can be accompanied by altered amplitudes after SD (Fig 5B, 104 resp. 62, dampened resp. increased amplitudes), meaning that the contribution of the C component to gene expression dynamics, relative to that of the S component which is constant, is modulated after SD. For example, for *Dbp*, breaking down the contribution of the S and C_A_ part of the dynamics show that in baseline, the expression is influenced by both sleep-wake history and time of day, while after SD, the contribution of time of day is diminished by 90% and thus the recovery dynamics are driven mainly by S (Fig 5B, blue dotted curve). Other examples of genes with dampened amplitudes include *Per2* (Fig S2 page 18), a gene known to be subject to complex interactions between Processes S and C (28), as well as clock output genes *Nfil3*, and *Bhlhe41* (*E4bp4*, resp. *Dec2*; Fig S2 pages 19-20). S+C_A_ genes peak during the first half of the light, resp. dark phases, at yet a different time than all other models. Still, genes displaying decreased expression under waking had phases overlapping with model C_A_ (Fig 5A).

### Recovery time course uncovers hitherto unnoticed genes affected by SD

Strikingly, a majority of the genes assigned to the amplitude-affected models C_A_ and S+C_A_ (759 out of 960 genes, 79%) were not identified when we examined differential expression at the end of SD alone (*i.e.* T30 vs. T6), as in previous studies (e.g. (13, 21, 29)). For example, fatty acid binding protein 7, *Fabp7* (model C_A_, Fig S1A, top left), is not differentially expressed at T27 nor T30, however its oscillation amplitude displays the strongest reduction from T36 onwards among non-DE genes, possibly due to its being a target of *Nr1d1* (model C_A_, differentially expressed at T27 and T30) and thus downstream of the primary response (30). This was especially true for the C_A_ model, where only 55 out of 794 (7%) genes were differentially expressed at ZT6, vs. 118 out of 166 (73%) for the S+C_A_ model (as a comparison, 1746/2677 (67%) genes in the S model were significantly differentially expressed at T30, and 468/2457 (6.5%) genes in the C model). This observation highlights the importance of examining the molecular recovery process from SD over time, considering dynamics due to both spontaneous and enforced wakefulness in the same experiment.

Conversely, genes that were differentially expressed at T27 or T30 (2862 genes) were enriched in models S, S+C and S+C_A_, and underrepresented in models C and C_A_ (Fig 4F). Interestingly, 375 of these genes were assigned to model F, representing genes acutely affected by SD, which is not equivalent to sleep-wake driven, as their overall time course is perturbed only at one or both of these SD time points, but otherwise not modulated by the sleep-wake distribution.

To assess how expression patterns return to baseline, we examined differential expression at each time point during and after SD compared to baseline (*i.e.* T27/T3, T30/T6, T36/T12 etc. until T78/T6) and found 210 genes genome wide that were differentially expressed after phenotypic recovery (*i.e.* after T42), namely 137 genes at T48 and 75 genes at T60. This was consistent with the observation that the proportion of genes with a *p*-value <0.05 in the cluster analysis did not reach zero for all post-SD time points (Fig 2C). Also, because model C_A_ (as well as model S+C_A_) is penalized for complexity, the change in amplitude needs to be pronounced and long-lasting for a gene to be assigned to it. Thus, the assignment of any genes to models C_A_ and S+C_A_ implies that SD alters amplitudes in gene expression rhythms and generally affects gene expression dynamics beyond SD. Taken together, these observations show that the molecular perturbations outlast the phenotypic changes, meaning that the mice have not yet recovered from SD despite behavioral and electrophysiological measures of sleep need having returned to baseline.

### Genome-wide ATAC-seq analysis shows a rapid response and sleep-wake driven dynamics in chromatin accessibility

We next asked which regulatory elements are underlying the extensive transcriptome response to SD. We used ATAC-seq (31) to identify a union of 130’727 chromatin accessible regions (called peaks, see Methods) over all time points (*i.e.* a peak is present in at least one time point). The majority of detected peaks do not change over time and are constantly accessible. Indeed, while 25% of expressed genes show a time dynamic according to a likelihood ratio test at a 0.001 FDR threshold, only 3.7% of ATAC-seq peaks display a time dynamic at the same stringent threshold. While the first principal component (PC1, 10%) probably represents experimental noise, 7% of the variance among time points, represented by PC2, could be attributed to the sleep-wake history and follows sleep-wake dynamics, paralleling the RNA-seq data (Fig 6A, PC2 and 6B). The accessibility of these regions mimics an IEG response to SD with a very fast modulation of the chromatin, detected already after the first 3 hours of SD, a striking illustration of the plasticity of this compartment.

**Fig 6.**
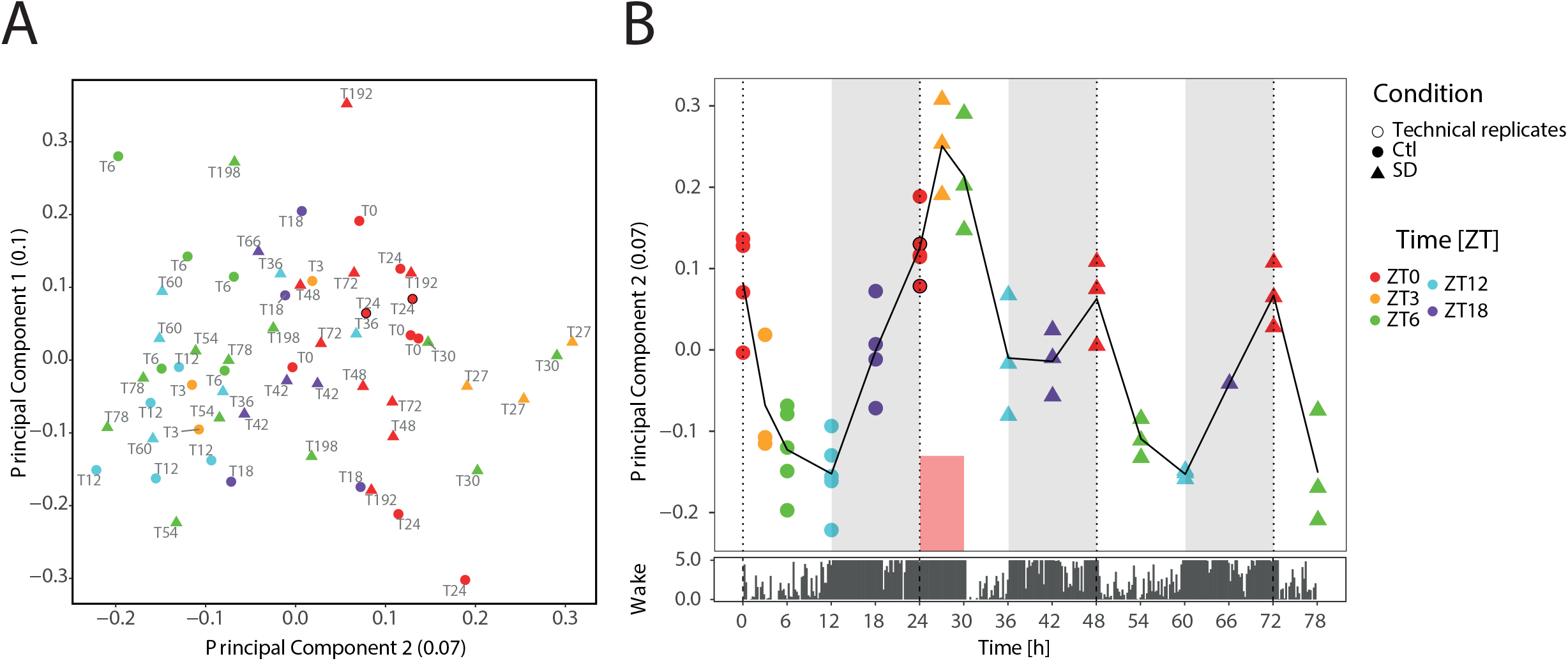
Chromatin accessibility shows sleep-wake driven dynamics and a rapid response to SD. **(A)** Principal component analysis of all detected ATAC-seq peaks. Plot features as in Fig 2A. A black symbol outline highlights technical replicates. **(B)** Second principal component plotted over time. Color and shape code as in A.

The strongest differential signal relative to baseline occurred during SD with 1793 peaks differentially accessible (differentially accessible sites, DAS) at ZT3 (T27 vs. T3, after 3h SD); 2098 at ZT6 (T30 vs. T6, end of 6h SD), with 607 peaks in common. Differential signal during SD (ZT3 and ZT6) consisted predominantly of increased accessibility (91% of DAS for ZT3 and 88% of DAS for ZT6), while the 645 late-responding DAS (*i.e.* differentially accessible at ZT12 only, T36 vs. T12, after 6h of recovery, Fig S3A) were more likely to be decreased (55.5%).

To verify that the DAS we identified followed the homeostatic process, we asked whether the effect of spontaneous waking during the first 6 hours of the baseline dark phase (ZT12 to −18, when mice are predominantly awake; see Fig 1B) was similar to the effect of forced wakefulness (SD). We therefore tested, for each of the 2098 DAS induced by SD, whether we could reject the null hypothesis of an identical fold-change induced by SD (T30 vs. T24) from the fold-change in baseline (T18 vs. T12), and found that only for 296 DAS (17%) the null hypothesis had to be rejected (uncorrected *p*-value < 0.05). Therefore, the majority of the DAS we identified display a similar response to spontaneous and forced wakefulness, and are thus likely to be sleep-wake driven instead of being affected by other factors associated with the SD protocol such as stress.

Overall, dynamics in chromatin accessibility were most pronounced during SD, and no longer significantly differed from baseline by 12 hours after SD (T42). Although we cannot exclude that this seemingly faster recovery is due to a lower sensitivity of the ATAC-seq signal relative to RNA-seq, these results do show that changes in chromatin accessibility start appearing early in the response to SD, confirming that chromatin accessibility is dynamic and can change on short time scales, even faster than observed in circadian oscillations (32).

Consistent with previous studies (e.g. (33–35)), accessibility peaks from all time points and conditions were mainly located in intronic or intergenic regions (Fig S3B). When considering only DAS sites (at ZT3, −6, or −12), the proportion of intergenic regions was increased at the expense of the other regions (Fisher’s exact test FDR adjusted *p*-value <0.01), suggesting that SD influences the accessibility of distal rather than proximal elements (Fig S3C-E). Genes associated with DAS (see Methods) were enriched among models involving sleep-wake driven dynamics (Fig S3F-K) compared to all peaks, for DAS dynamic groups at ZT3, ZT6 and combinations of ZT3, −6 and −12 (see above, Fig S3A, *p*-values < 2e-10, chi-square test), but not at ZT12 only (*p*-value = 0.48).

### Gene expression correlates with chromatin accessibility at distal elements rather than promoters

Probing the general dynamics of chromatin accessibility by k-means clustering, we found three main types of temporal profiles, all of which were reminiscent of sleep-wake driven dynamics (Fig S3L). Clusters 1-3 displayed an early response to SD with a fast recovery, clusters 4-7 also present an immediate response, but a slow recovery, while clusters 8-10 displayed a late response. Because of this similarity with the RNA-seq clustering (RNA clusters 1-8), and because of the similarity in the general dynamics observed by PCA between RNA-seq sand ATAC-seq, we sought to connect these changes in accessibility with changes in gene expression and, as both signals originate from the same mouse, correlated ATAC-seq peak signal over time to gene expression levels over time by calculating the Pearson correlation across samples (Methods). We confined the possible peak-to-gene associations to a single ATAC-peak per gene within the same topologically associated domain (TAD) defined from Hi-C data generated from mouse cerebral cortex (36). In total, the expression level of 3294 genes was significantly associated with the ATAC-seq signal of one peak, at distances ranging from the transcription start site (TSS) to 5 Mb away (mean distance for all significant associations at 0.05 FDR: 0.65 Mb). We observed both positive and negative correlations between expression and ATAC-seq signal, implying that both enhancers and repressors are involved in the response to SD (Fig 7A).

**Fig 7.**
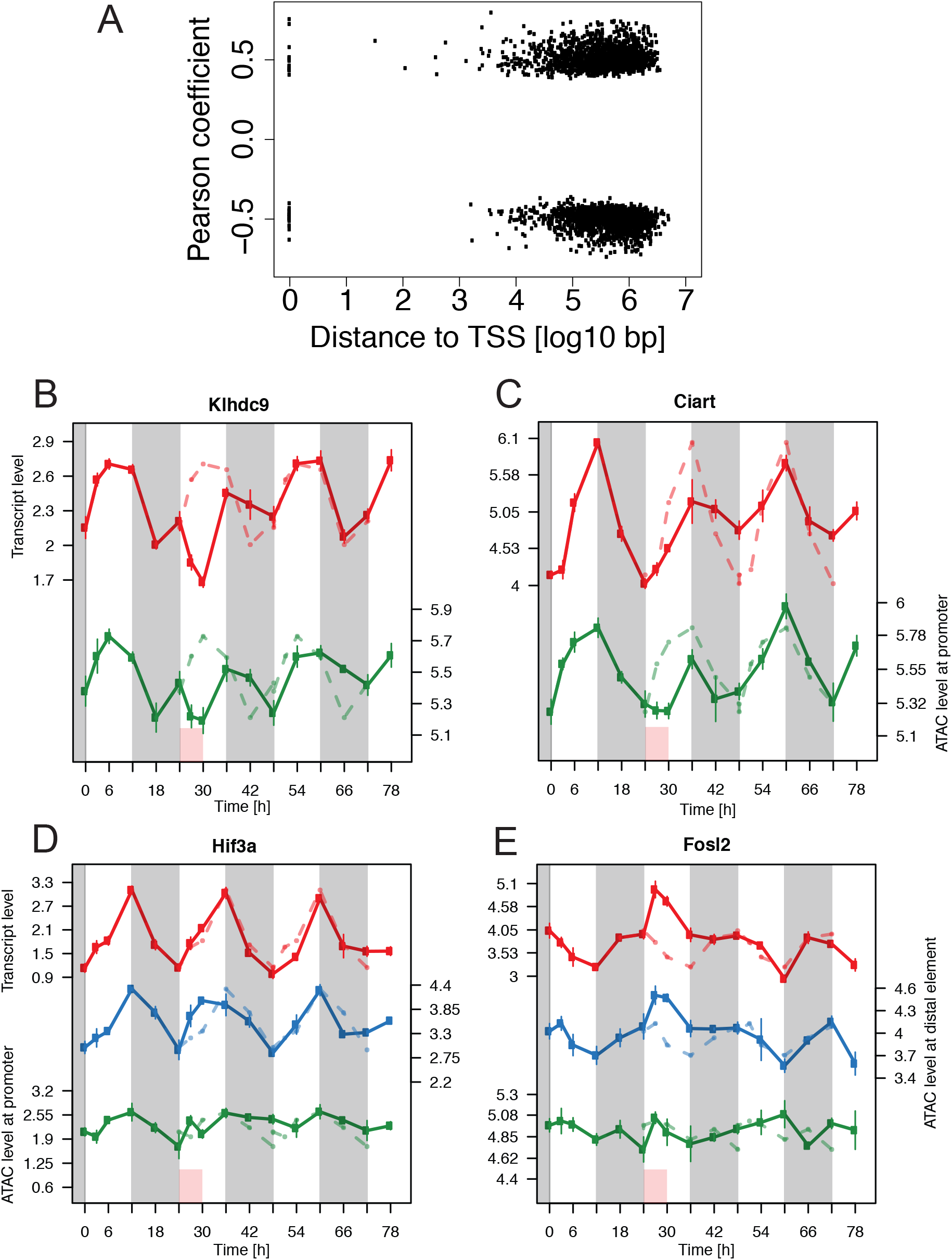
Gene expression predominantly correlates with the dynamics of distal accessible genomic regions rather than promoters. **(A)** Distance from ATAC-seq peak to the associated TSS. **(B-E)** Temporal patterns of gene-peak associations (with promoter for genes associated with distal elements). Top: RNAseq signal. Middle and bottom: ATAC signal of the distal element, resp. promoter, of the associated gene. Red box: SD. Grey shading: dark phase of the light-dark cycle.

Among the strongest associations (|ρ|>0.7, 34 pairs, Table S5), we found mostly genes assigned to models involving sleep-wake driven dynamics (*i.e.* models S, S+C, S+C_A_, respectively 26, 4, 1 gene), but also two genes from models C and C_A_ (*Hif3a*, resp. *Gm13889*) and one gene from model F (*Mid1*). Only 38/3294 significant gene-peak associations were proximal (*i.e.* the peak spanned the TSS), overlapping with only two genes of the top 34 pairs, namely *Klhdc9*, an interaction partner of CDK2-associated cyclin A1, *Ccna1* (37) (model S, Fig 7B), and *Ciart* (Model S+C_A_, Fig 7C), an interaction partner and suppressor of the Arntl and Per2 proteins (38), thereby forming an additional negative feedback loop to the circadian molecular machinery. Also, because *Ciart* is sensitive to and affects stress signaling pathways (38, 39), it can be conjectured that the stress associated with SD (21) could be a factor contributing to the sustained circadian dysregulation of the clock gene circuitry we have found in the cortex.

Thus, the majority of genes correlated more strongly with a distal element than with the accessibility of their promoters. Generally, the mean |ρ| value of the correlation between the expression and the ATAC signal spanning the TSS of all genes involved in an association was 0.16, while it was 0.5 for the correlation between expression and the ATAC signal of the top associated distal peak for the same genes. The top correlations to a distal element involved the environmental sensor *Hif3a* (model C, Fig 7D), and the immediate early gene *Fosl2* (model S, Fig 7E). In the case of *Hif3a*, the distal element (blue line) displayed an immediate response to SD with an increase at T27 and T30, possibly a relative decrease at T36 before resuming the baseline pattern from T42. The promoter (green line) likewise showed a response at T6 followed by a slow recovery from T36 to T54, while the RNA oscillation (red line) was largely unperturbed. For *Fosl2*, we observed a fast response of the distal element together with the mRNA, plateauing already at T27 and followed by a fast recovery by T36, whereas the promoter followed a different pattern. We note that the variability of the ATAC-seq signal can hamper the exact definition of the dynamics of promoters and distal elements. We observed similar relationships between gene expression and accessibility of the corresponding promoter and associated distal peak for the remaining genes of the top 34 correlations (Fig S4).

The widespread lack of correspondence between promoter and transcription dynamics hint at a model where transcription happens from an accessible promoter under the regulation of a distal element mediated by transcription factors (TF). A TF motif activity analysis (40, 41) taking advantage of our paired RNA-seq and ATAC-seq data predicted the SRF (serum response factor) motif to be by far the most statistically significant candidate in the entire temporal gene expression dataset (*i.e.* the genes assigned to models S, C, C_A_, S+C and S+C_A_, Fig 8A). The inferred temporal activity of SRF (Methods) was consistent with sleep-wake driven dynamics, paralleling the expression of the *Srf* transcript (Fig 8B-C). The genes with the strongest contribution to the enrichment signal, namely *Egr2*, *Junb, Fos, Arc*, and *Nr4a1*, are immediate early genes and were all classified under model S, as was *Srf* itself (Model S, *w*=0.979). Scanning the open chromatin regions corresponding to <5 kb up- and downstream of the promoters of these genes, we found SRF binding sites which overlapped with ChIP-seq peaks against SRF in mouse fibroblasts (42) (see *Egr2* as example in Fig S5). Thus, the correlation of gene expression with the accessibility of distal elements rather than their promoters, together with the presence of SRF motifs, suggests a model of the response to extended waking where SRF is bound to a constitutively open promoter, ready for an interaction with a distal element that changes its own activity and mediates the changes in gene expression. We note that while *Srf* was not identified as DE after SD in previous studies, we found it to be more strongly DE after 3h than after 6h SD. This observation, along with the fast increase in chromatin accessibility, highlights the importance of increased time resolution in sampling, particularly in the early hours of SD.

**Fig 8.**
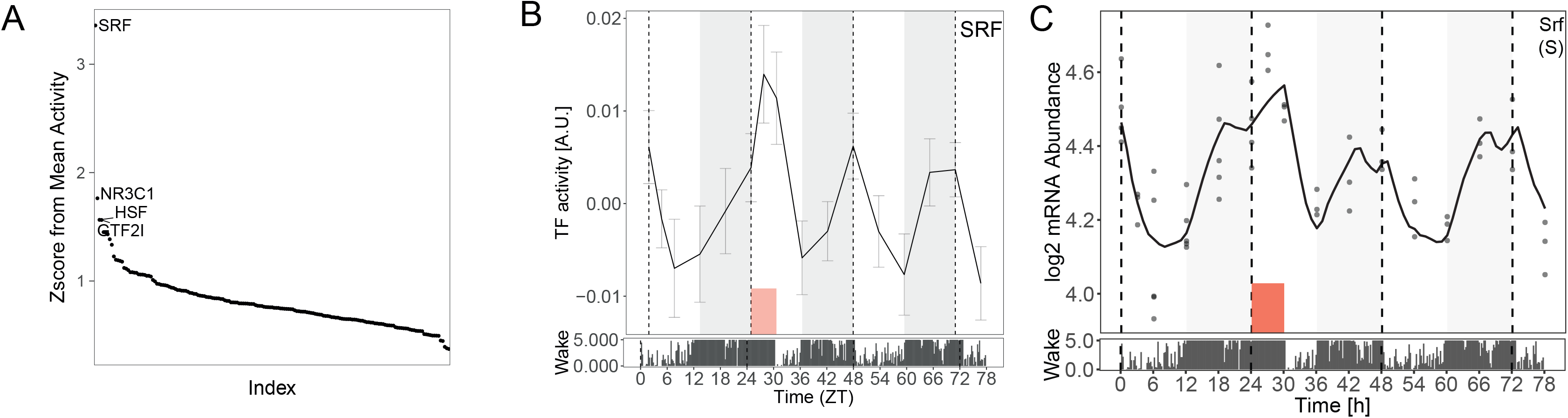
SRF activity in the response to SD. **(A)** 179 TF motifs ranked by z-score that explains the temporal dynamics in the RNA-seq dataset. **(B)** Inferred temporal activity of SRF. Error bars are standard deviations of the activity estimates. Lower bar denotes time spent awake in 5-minute bins. **(C)** *Srf* transcript time course.

## Discussion

We have characterized the dynamics in cerebral cortex of transcriptome and regulatory elements in relation to the sleep-wake distribution before, during and after an exposure to sleep deprivation, a one-time and short intervention during the first half of the habitual rest phase of mice. We found that changes in both transcriptome and chromatin accessibility are largely governed by a sleep-wake driven process with or without interaction with the circadian process, and that the molecular recovery from SD outlasts the electrophysiological and behavioral signs of sleep need.

We developed a model selection approach integrating sleep-wake states to classify genes according to sleep-wake driven dynamics, time-of-day dynamics, interactions between the two, with the possibility of an alteration of the oscillation amplitude. This set of models allowed us to classify the genes according to their temporal expression pattern, and determine the relative contributions of and interaction between the circadian and sleep-wake processes governing the expression of genes with 24-hour rhythms. This classification proved more powerful in identifying sleep-wake driven genes than past single-time-point differential expression studies. Further studies using higher sampling resolution for a longer time will allow to apply a more refined statistical framework to the recovery phase in particular and thus further classify the response and recovery patterns.

A striking new finding was that most clock genes were affected by SD. Previous work had already shown that the cortical expression of a number of clock genes is affected during SD (for reviews see (29, 43, 44)) and that SD acutely suppresses the specific DNA-binding of the circadian transcription factors BMAL1 and NPAS2 to their target genes *Per2* and *Dbp* (45), demonstrating that prolonged wakefulness intervenes directly at the core of the circadian molecular machinery. An important long-term consequence of this intervention could be the dampened amplitude we observed in the rhythmic expression of most of the known clock genes (i.e., *Arntl*, *Per2*, *Per3*, *Nr1d1*, *Nr1d2*, *Cry2*, *Ciart*, and the clock-output genes *Dbp*, *Nfil3*, *Bhlhe41*, all of which were model C_A_ and S+C_A_ genes). This long-term dampening of clock gene rhythms was all the more surprising given the fact that the observations were made under entrained light-dark conditions and in the presence of a largely unaffected diurnal sleep-wake distribution, two factors known to contribute to high amplitude clock gene expression. Because disrupted clock gene rhythms have been causally implicated in the etiology of disease like metabolic syndrome (reviewed in (4)), clock genes could be a final common molecular pathway underlying the etiology of metabolic syndrome associated both with insufficient good quality sleep and with circadian misalignment (43).

Studying chromatin accessibility for the first time in sleep research allowed us to identify a set of genomic regions as first actors in a possible repertoire set in motion already after 3h SD and giving rise to differential gene expression. The increased proportion of distal elements among DAS compared to all regions, together with the correlation of dynamic gene expression with distal elements rather than the respective promoters is consistent with a scenario where expression is modulated by different enhancers or repressors interacting with an accessible promoter under the influence of regulator proteins. The implication of SRF as a candidate priming factor in the response to SD is compelling, as it plays a key role in activity-dependent modulation of synaptic strength (46), and its ortholog *blistered* is required to increase sleep after social enrichment in *Drosophila* (47, 48).

Our results imply that beyond an apparent recovery from SD lie deeper, complex and longer-lasting molecular perturbations, even among clock genes. We also show that genes can seem unchanged when sampled at a single time point after SD, yet be affected by a profound perturbation later on. These perturbations eventually recover, as hinted by the absence of differential expression after 7 days, however until baseline is reached, this temporary regulatory background could possibly cause the response to another exposure (repeated SD or other) to differ from that under the pre-SD baseline background. While it is debated whether repeated sleep deprivation on subsequent days alter the homeostatic response at the phenotypic level in rats (EEG) (49, 50), recent studies in humans found that even two nights of recovery sleep were insufficient to completely reverse the metabolic perturbations caused by multiple nights of restricted sleep (51, 52). Follow-up experiments at the molecular level will show how such a transient “new baseline” due to partial recovery would influence the response to a second event occurring before full recovery.

## Materials and Methods

### Animals

C57BL/6J male mice were purchased from Charles River France (Lyon, France) and allowed to acclimate to our sleep study facility for 2-4 weeks prior to habituation to the experimental setting. Animals were kept in accordance to the Swiss Animal Protection Act, and all experimental procedures were approved by the local veterinary authorities.

### Surgery and EEG recording

The EEG cohort consisted of 12 male C57BL/6J mice 10-12 weeks at the time of SD that were part of another study (12). Surgical implantation of electrodes, EEG recording and data collection were performed according to our standard procedure (53). EEG was recorded from 2 days prior to SD (which were averaged to constitute a 24-hour baseline) until 2 days after SD. In a subset (6/12) an additional 5 days were recorded. Electrophysiological signals were captured and transformed from analog to digital with a sampling rate of 2000Hz, and down-sampled and stored at 200Hz (*EMBLA A10* and *Somnologica-3*; Medcare Flaga; Thornton). Sleep and wake states were annotated according to established criteria based on the properties of the EEG and EMG signals (53). To determine spectral composition, EEG signals (0 to 90 Hz) underwent a discrete Fourier transformation, using a window of 4 seconds (Hamming function), to determine power spectral density. Delta power (1-4Hz) was extracted for NREM sleep epochs, averaged over consecutive intervals to which an equal number of 4-second NREM sleep epochs contribute (*i.e.* percentiles), and then expressed as a percentage of the levels reached between ZT8-12 (when both delta power and sleep homeostatic pressure reach lowest levels during baseline) during the 2 baseline days (see (19) for details). SD and recovery time points were compared to baseline by means of 2-way repeated measures ANOVA followed by post-hoc t-tests.

### Sleep deprivation and tissue collection

Mice for tissue collection were divided into two experimental cohorts, sleep deprived (SD) and non-sleep deprived (controls, Ctr). After a one-week habituation to the experimental setting, at the age of 11-12 weeks, the SD mice were sleep-deprived by gentle handling for 6 hours starting at light onset (zeitgeber time ZT0-ZT6) as described in (53), and allowed to recover according to the tissue collection schedule. Mice were anesthetized with isoflurane prior to decapitation. Cortex was rapidly dissected and flash frozen in liquid nitrogen. Below, we refer to each time point in hours from the start of the baseline day (T0) until the end of tissue collection on Day 8 (T198), with SD occurring from T24 to T30. The study design is represented in Fig 1A: control mice were sacrificed at ZT0, ZT3, ZT6, ZT12 and ZT18 of the first day of experimentation (samples T0-T18), serving as a baseline day (Day 0). On Day 1, SD mice were sacrificed at the same time of day as on Day 0 (samples T24-T42, with T27 and T30 samples being taken after 3h and 6h SD, respectively), on Day 2 at ZT0, ZT6, ZT12, ZT18 (samples T48-66), as well as ZT0 and ZT6 on Day 3 (samples T72-78). Sampling at ZT3 on Day 0 and Day 1 served to provide an intermediary time point increasing the time resolution during SD. Finally, two groups of mice were allowed to recover for 7 days after SD, before being sacrificed at ZT0 and ZT6 (Day 8, samples T192-198). We collected 3-4 replicates per time point and condition, and 8 replicates of ZT0 controls from two different animal batches, which were divided evenly between T0 and T24 in the analysis. T192 and T198 were collected to probe the persistence of the effects detected during Days 1-3. The clustering and model fitting analyses used time points T0-T78.

### Tissue processing and sequencing library preparation

Frozen cortex of each individual was ground in liquid nitrogen and stored at - 80°C until further use. Tissue from each mouse was distributed to the two protocols (RNAseq and ATAC-seq), such that both datasets originate from the exact same set of individuals, allowing us to use the paired information when correlating the two datasets (see below). The only exception was time point T66, where two out of three ATAC-seq replicates needed to be excluded from the analysis due to sequencing failure.

Total RNA was extracted using the miRNeasy kit (Qiagen; Hilden, Germany) following the manufacturer’s instructions.

RNA-seq libraries were prepared using 1000 ng of total RNA and the Illumina TruSeq Stranded mRNA reagents (Illumina; San Diego, CA, USA) on a Sciclone liquid handling robot (PerkinElmer; Waltham, MA, USA) using a PerkinElmer-developed automated script. Libraries were sequenced on the Illumina HiSeq 2500 sequencer, producing >36 million (median 55 million) mappable single-end 100 bp reads.

ATAC-seq was performed with minor modifications from (54). 100’000 nuclei were treated with 2.5 µl Tagment DNA enzyme (Nextera DNA Sample Preparation Kit, Illumina) in transposition buffer (10mM Tris Base, 5mM MgCl_2_, 10% DMSO, pH 7.6, adapted from (55)) at 37°C for 30 minutes, followed by cleanup on a Qiagen Minelute column. Fragments >1kb in size were removed using 0.6X, then 1X, volumes of AmpureXP beads (Beckman Coulter Life Sciences; Indianapolis, IN, USA). DNA fragments were subjected to 11 cycles of PCR amplification with Nextera dual index primers (Illumina) and the NEBNext High Fidelity 2X PCR Master Mix (New England Biolabs; Ipswich, MA, USA). PCR reactions were cleaned up with one volume AmpureXP beads, quantified by Qubit (ThermoFisher Scientific; Waltham, MA, USA) and quality controlled by Fragment Analyzer (Advanced Analytical Technologies; Ankeny, IA, USA). Libraries were sequenced on the Illumina HiSeq 2500 sequencer, producing >25 million (median 41 million) mappable 50 bp paired-end reads per sample after removal of duplicate and mitochondrial sequences.

### Sequencing data analysis

Transcript abundance was quantified by *kallisto* version 0.43.0 (56) using the GRCm38 reference transcriptome (mm10) and the parameters `--single -l 100 -s 20 -b 100`. The abundances were processed as follows using *sleuth* version 0.29.0 (57): transcript abundances were merged into gene counts in transcripts per million (TPM), after which we applied a detection cutoff of 5.5 on the mean gene counts across samples in the time series, yielding a set of 13’842 expressed genes which were used for further analysis. Batch effects were corrected by *ComBat* (R package *sva*_version 3.25.4 (58)). Batch-corrected transcript abundances and scaled abundances are given in Tables S1 and S2. For genome browser visualization, sequence reads were aligned to the mouse genome (mm10) using *kallisto* version 0.44 with the same alignment parameters used for quantification and transformed into bam files using the `–genomebam` parameter with the *Mus_musculus.GRCm38.93.gtf ensembl release 93* annotation file. Alignment files were finally converted to bigwig using *deepTools* (59).

ATAC-seq reads were aligned to the mouse genome (mm10) using *bowtie2* (60) in paired-end mode, with the parameters recommended for open chromatin (`-- very-sensitive --maxins 2000 --no-mixed --no-discordant`). Duplicate sequences were removed using *samtools* rmdup (61).

### Differential gene expression

Differential expression at each time point was performed using the Wald test, implemented in *sleuth* version 0.29.0 (Pimentel et al. 2017). Each time point during and after SD was compared to the corresponding baseline time, *i.e.* the same ZT time. We note that expression levels at T192 and T198 were not significantly different from baseline at T0, respectively T6 (FDR adjusted *p*-value > 0.05).

### Clustering of mRNA profiles

To uncover temporal patterns of mRNA abundance, we performed *k*-means clustering on genes displaying statistically significant temporal expression, defined as follows: to identify genes displaying a statistically significant effect over time, we used a likelihood ratio test implemented by *sleuth* version 0.29.0 (57), comparing a full model with a parameter for each time point plus a batch effect (*i.e.* t=[0, 3, 6, 12, 18, 24, 27, 30, 36, 42, 48, 54, 60, 66, 72, 78] plus a batch effect) versus a null model with no time effect (*i.e.* only a batch effect). We used an FDR-adjusted *p*-value cutoff of 0.001, which yielded 3461 statistically significant genes, which were used in the clustering analysis. This conservative cutoff was adopted to ensure the discovery of robust temporal patterns. For a range of number of clusters, *k*, we calculated the within cluster variation as the sum of the Euclidean distance between data points and their assigned cluster centroids and empirically chose *k*=10 as a balance between variance explained and generalizability of each cluster. The proportion of genes at each time point with a *p*-value < 0.05, as calculated from a likelihood ratio test between SD and Ctr, is represented by a grey shaded bar above each cluster.

### mRNA time course analysis

We used a model selection approach to classify the temporal log mRNA abundance *m*(*t*) of all 13’842 expressed genes into the scenarios described in Results and represented in Fig S1B. The models can be expressed as stated below. For models 2, 5 and 6, sleep-wake history was used to model the synthesis rate of mRNA according to Process S in the 2-two process model (19) using sleep-wake data from n=12 C57BL/6J mice of the same age and sex and recorded under the same conditions (12).

1. Flat model with constant *μ* and noise *ϵ* (F)

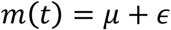
2. Sleep-wake model (S)

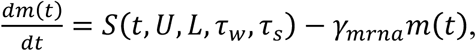

where the production function S is defined recursively:

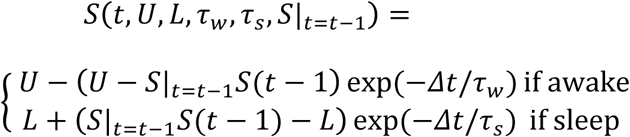

with Δt: mean period of continuous wake or sleep, defined by sleep-wake data from 12 mice *U*: asymptotic value for long periods of wake *L*: asymptotic value for long periods of sleep *γ*_*mrna*_: degradation rate of mRNA *S*|_*t=t-1*_: previous value of S *S*|_*t=0*_ = *S*_*0*_: initial value of S

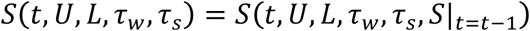 The interpretation of this model is that mRNA abundances are driven by regulatory dynamics that follow Process S. Including a degradation rate of mRNA γ_*mrna*_ allows genes driven by the sleep-wake distribution but having long half-lives to still be fit by the sleep-wake model, since a delay in the response is then expected. We solved the differential equation for *m*(*t*) using the Euler method with a time step of 0.1 hours. We will call the solution of this differential equation 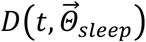 where 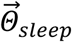 are the sleep parameters, *S*_O_, *U*, *L*, *τ*_*w*_, *τ*_*s*_, *γ*_*mrna*_. The model we try to fit is therefore:

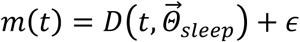
3. Cosine oscillatory model (C)

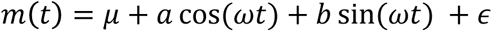

with: *μ*: mean of signal √(*a* + *b*)^2^ mean to peak amplitude of signal tan^−1^ (*b/a*): phase of the signal *ω* = 2Π/24: angular frequency ϵ: Gaussian noise
4. Cosine model with change in amplitude (C_A_).

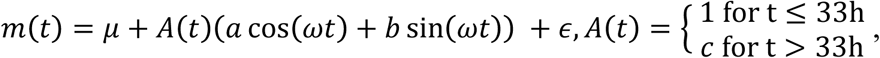

where t=33 h corresponds to 3 h after the end of SD. Thus, in this model the amplitude is changed by a factor *c* after *t* = 33 h, *i.e.* between T30 and T36. We have chosen to allow for a single change over the time course, as the time resolution and sampling length in time only allows to confidently follow one complete oscillation cycle.
5. Sleep-wake and oscillatory model (S+C)

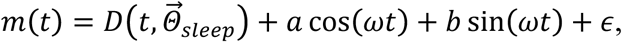

where 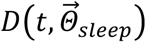 is the solution to the differential equation in the sleep model.
6. Combined with change in amplitude model (S+C_A_)

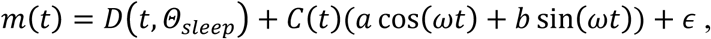

where D and C are defined as above.

Of note, we also included a generic model:

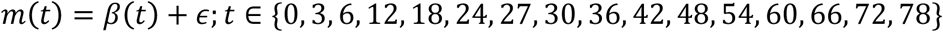

where the gene expression is modeled as the mean expression across replicates at each time point to assess the possibility of more complex dynamics not explained by any of the 6 models. We found that the BIC weight *w* was always lower for this model than the other 6, meaning no genes were assigned to it, and therefore not included in subsequent analyses.

For models that are nonlinear with respect to the parameters (models 2, 4-6), we fitted the model with the optim() function in R using the L-BFGS-B method. To constrain time constants in the S process such that resulting predictions are at steady state during baseline, we penalized the negative log likelihood by 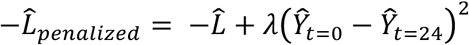, where *λ* = 1000 is a penalization parameter, 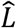 is the log-likelihood from the fit, and 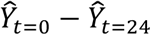 is the predicted log gene expression difference at t=0 and t=24, respectively. This penalizes predictions that deviate from steady state in baseline. Linear models (models 1, 3, and generic) were solved using the lm() function in R. The mRNA levels were fit in the log scale.

For each gene, we estimated the posterior probability of each model by first calculating the Bayesian Information Criterion (BIC) scores:

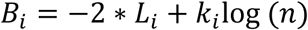

where *L* is the log likelihood. A better fit will improve (decrease) the BIC, while a more complex model will penalize (increase) the BIC. Intuitively, an optimal model will fit the data while not using an excessive number of parameters. We assume the model errors are independent and identically distributed following a Gaussian distribution with variance estimated from the fits:

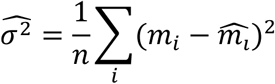

Exponentiating the BIC scores yields Schwarz weights *w*_*i*_:

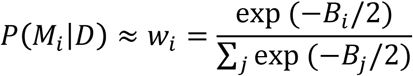

We then assigned each gene to the model *i* corresponding to the largest *w*_i_.

*w*_*i*_ assigns a probability to each model, and this probability measurement takes into account the number of parameters *k* in the model through the BIC score (*i.e.* complex models with large *k* are penalized by having a larger *B*, which would have smaller *w*). All genes were assigned to one model, 11141/13842 (80.5%) with a *w*>=0.6, and 12111/13842 (88%) with a difference >=0.2 to the second ranking *w*.

### Harmonic regression in baseline

To detect genes with rhythmic expression in baseline, we used harmonic regression on the Day 0 time points, which were fit using a linear model:

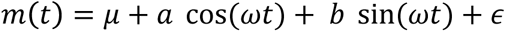

Where:

*μ*: mean of signal

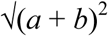 mean to peak amplitude of signal

tan^−1^(*b*/*a*): phase of the signal

*ω* = 2Π/24: angular frequency

ϵ: Gaussian noise

The parameters *μ*, *a*, *b* were fit using linear regression (lm() in R).

### ATAC-seq peak detection and quality control

ATAC-seq data files were processed before peak calling as follows. Alignment files were converted into bed files and tags were extracted using *bedtools* version 2.26.0. Each tag position was shifted +4 base pairs on the positive strand and - 5 base pairs on the negative strand to center tags on transposase binding events as suggested by (31). The peak calling was performed on pooled tags for replicates using *Macs2* version 2.1.1 (62) [`--nomodel --shift −75 --extsize 150`], and peaks were filtered using a 0.05 FDR cutoff for a local random Poisson distributed background noise, captured by *Macs2*. Peak boundaries were merged between time points and conditions in order to build a common peak mapping reference covering all samples, encompassing a total of 215’045 peaks. Finally, peak coverage was quantified using HTSeq version 0.6.1 for each sample, using the common mapping reference. We filtered low coverage peaks using a minimum mean threshold of 10 reads per peak and obtained 130’727 peaks.

We next performed two steps of quality control. First, we examined which genomic elements overlapped with our peaks and found that the proportion of ATAC peak basepairs mapping within introns and exons according to the *Ensembl_GRCm38/mm10 all genes* reference annotation (63) (62%) was higher than for the whole genome (44%), confirming that with ATAC-seq we are preferentially targeting active, *i.e.* accessible, parts of the genome. Second, we probed whether genes within accessible regions were enriched in cortex/brain tissue. To this end, we used the Bgee database and *topAnat* (64) to look for significant enrichment, and found that the top 20 enriched tissues were all nervous system structures (Table S3, FDR *p*-value < 10e-8). Finally, the proximity in the PCA of the two technical replicates at T24 attests the reproducibility of ATAC-seq over different batches of sequencing (Fig 6A).

### ATAC-seq clustering and differential accessibility analysis

To identify patterns of chromatin accessibility over time, we performed a clustering analysis using the same strategy as for gene expression. We identified 4824 sequences displaying a significant effect over time (LRT implemented in *edgeR*, FDR cutoff 0.001) and performed a k-means clustering (*k*=10).

To identify peaks with differential accessibility, we first normalized count data using a TMM normalization, applied a 10 read count threshold, and used a likelihood ratio test implemented in *edgeR*. We compared chromatin accessibility of SD samples (T27-198) with the corresponding ZT during baseline (T0-18, see Fig 1A). Thus, for differential accessibility at ZT3, we compared T27 with T3, at ZT6, T30 and T6, etc. *p*-values were adjusted using the Benjamini & Hochberg (FDR) method (65).

### Genomic distribution of ATAC-seq peaks

The annotation of the detected ATAC-seq peaks was performed using *PAVIS* with the *Ensembl_GRCm38/mm10 all genes* reference annotation (63).

### Peak-to-gene expression association

To associate gene expression dynamics with chromatin accessibility dynamics, we used a Pearson correlation coefficient across the samples and confined the possible association tests to previously defined topological interaction domains (TADs), which were computed from cortex tissue in Ref. (36). The positions of TAD boundaries were originally detected using the mm9 reference genome, so we converted them to mm10 using *CrossMap* 0.2.6 (66). For association statistics, we used a strategy similar to that implemented within *FastQTL* (67). Specifically, the time series of each pair, consisting of a peak and a gene within the same TAD, were associated using the Pearson correlation coefficient. For each gene, only the top correlated peak was retained. To control for multiple associations within a TAD and adjust nominal *p*-values, we used 1000 permutations per gene and modeled the null distribution fitting a beta distribution. The parameters were estimated using a maximum likelihood approach (R/MASS::fitdistr). Finally, a genome-wide *p*-value adjustment was computed using a *q*-value procedure (R/qvalue). Of the 11143 genes mapping within a TAD, 3294 were associated to an ATAC-seq peak within the same TAD using a 0.05 FDR cutoff.

### Prediction of transcription factor (TF) binding site (TFBS) activity in promoters

We inferred TF activity, based on the presence of TF motifs within ATAC-seq positive regions and the abundance of the nearby transcript, assuming that an accessible region containing TF binding motifs will be bound by the corresponding TF and transcription will occur as a result. Specifically, we used position weight matrices (PWMs) of 179 mouse transcription factors (TFs) defined by SwissRegulon on mm9 (http://swissregulon.unibas.ch). For each of the 179 PWMs, we scanned 500 bp windows within 15 kb upstream and 15kb downstream of transcription start sites using *MotEvo* (68) to obtain a site count matrix for each motif. We retained only regions containing ATAC-seq counts greater than 0.1 RPM (reads per million mapped reads). The site count matrix of each motif was scaled across genes so that ranges in site counts were comparable across motifs. We inferred TF activity using the TF binding site predictions and the temporal mRNA abundance, using a penalized regression model (*MARA*) as previously described (40, 41) and using an L_2_ norm penalty for regularization (ridge regression). Prior to the regression, we mean-centered the input matrix of temporal mRNA abundances, standardized the columns of the site count matrix (each motif across genes), and excluded genes that were assigned to the flat model (F).

## Data and code accessibility

Raw read files (.fastq), RNA transcripts per million (TPM), ATAC-seq peak calls and quantification are publicly available in the GEO/SRA repository under ID [to be communicated]. Code to run the model selection analysis is publicly available, found at https://jakeyeung@bitbucket.org/jakeyeung/sleepdepanalysis.git.

## Acknowledgments

We thank Shanaz Diessler, Marieke Hoekstra, Konstantinos Kompotis, Dessislava Petrova and the Lausanne Genomics Technologies Facility for technical assistance. Computations and analyses were performed at Vital-IT (http://www.vital-it.ch). CNH and MJ were funded by a grant of the Swiss National Science Foundation (31003A_173182) to PF. JY benefited from the Natural Sciences and Engineering Research Council of Canada Postgraduate Studies Doctoral scholarship.

