## Supplemental Figures and Table S3 for "Simple and complex interactions between sleep-wake driven and circadian processes shape daily genome regulatory dynamics in the mouse"

### Figure S1

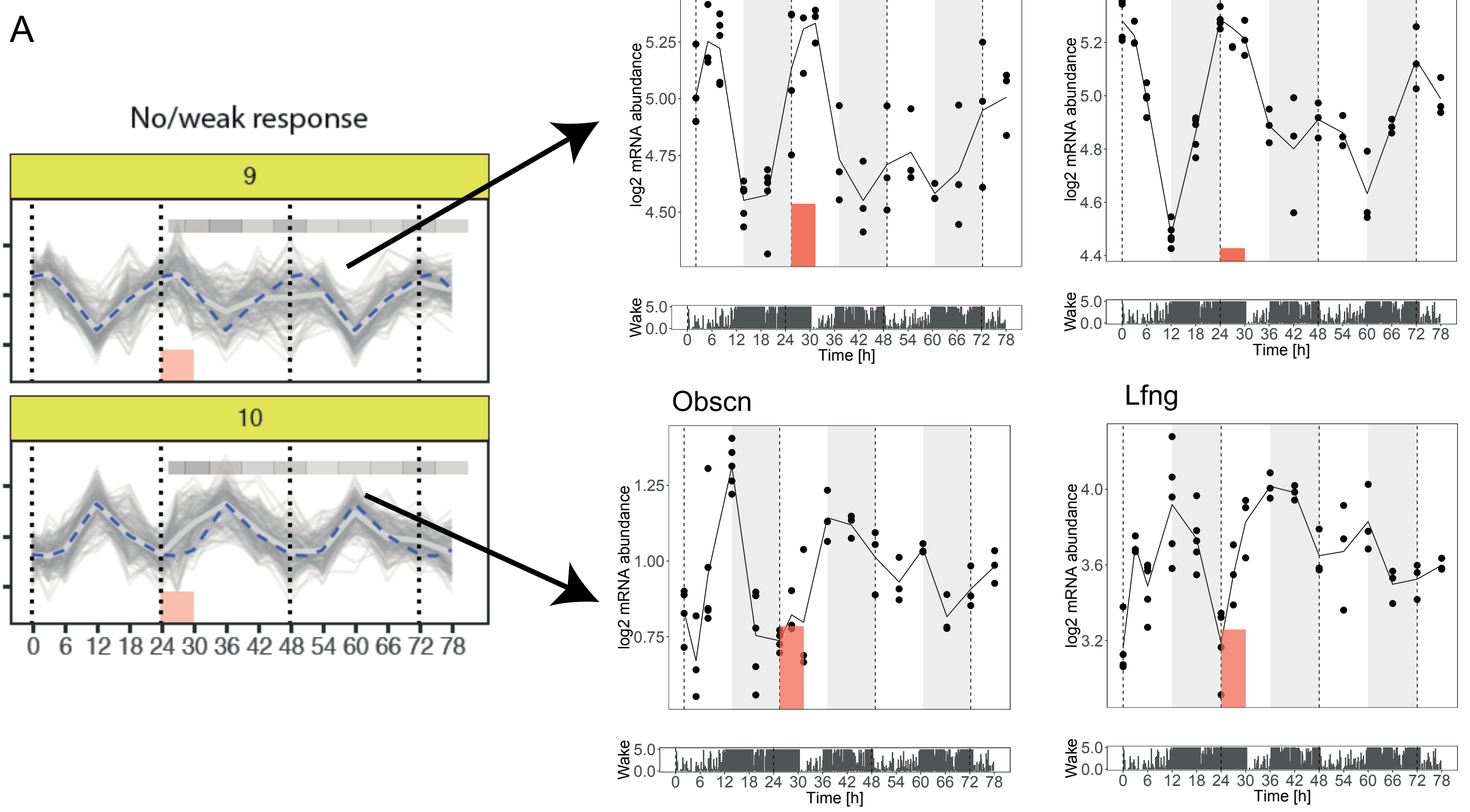


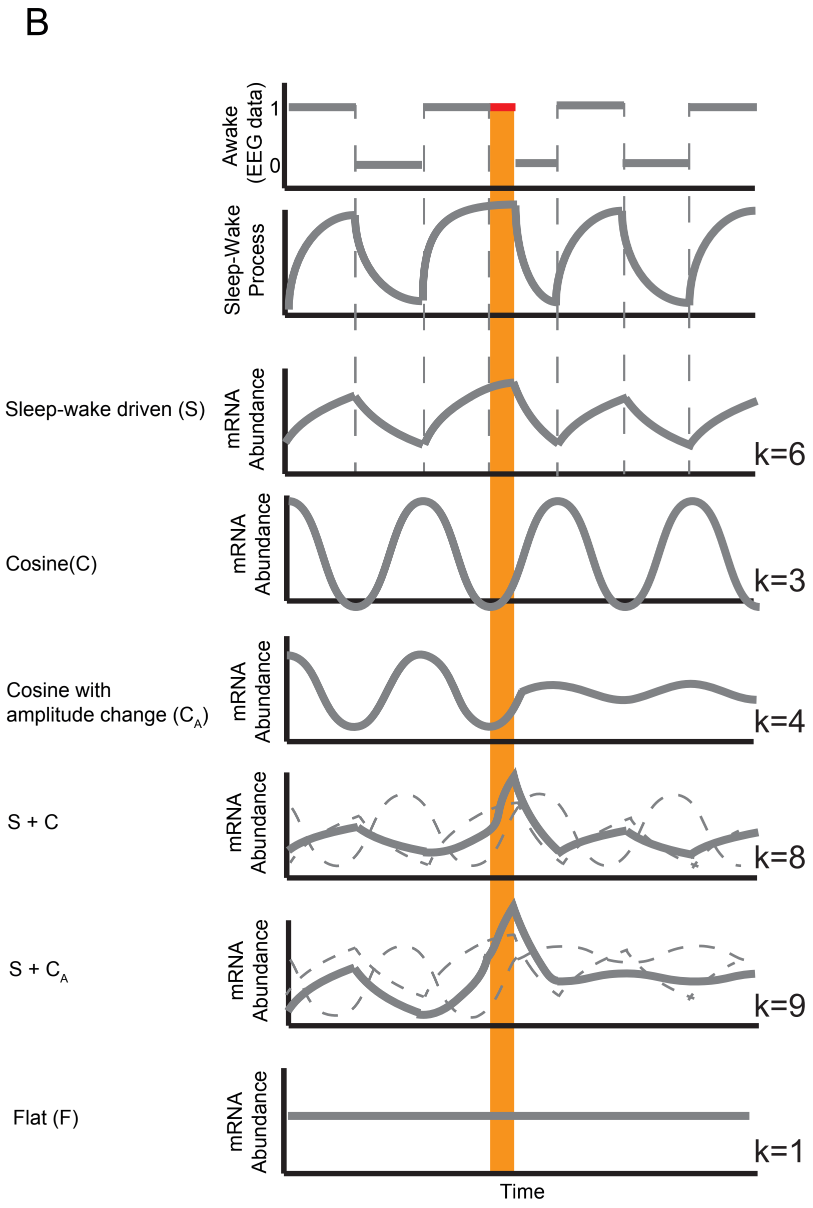


**Fig S1 Rationale for the model fitting**
**(A)** Examples of individual genes in cluster 9 and 10 with dampened amplitudes after SD. **(B)** Schematic of the 6 models for model fitting. Wakefulness state (top panel) is used to simulate Process S (second panel), which increases during wakefulness (awake=1) and decreases during sleep (awake=0). k is the number of parameters in each model. **(C)** Expression time course of Fabp7 as an example of an amplitude change without differential expression at the end of SD. Red box: SD. EEG data as in Fig 2.

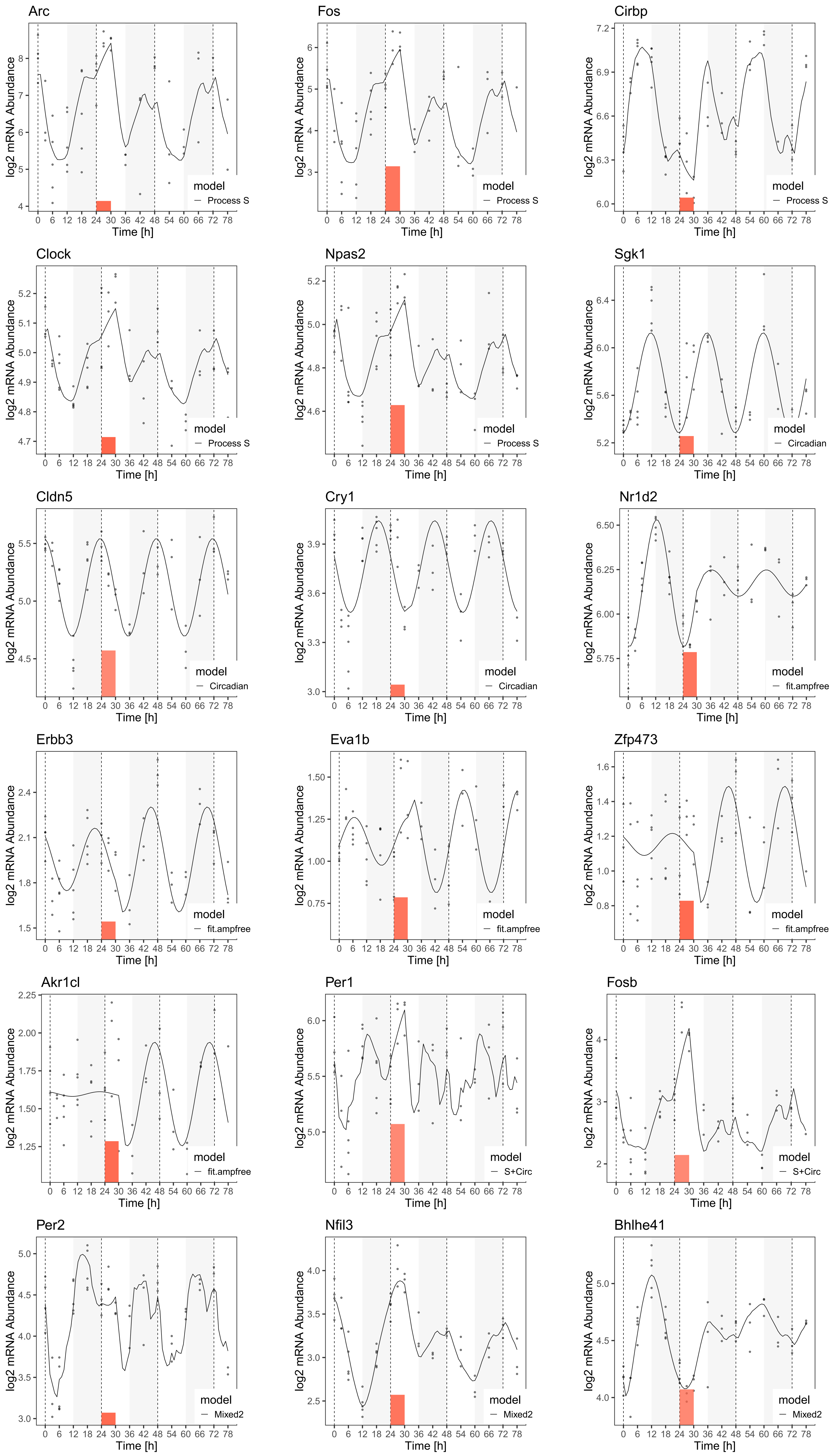


**Fig S2 Examples of invidual gene time courses discussed in the text**
Dots: RNA level data points. Bold line: best fitting model. Temporal EEG data as in Fig 2. Red box: SD.

### Figure S3

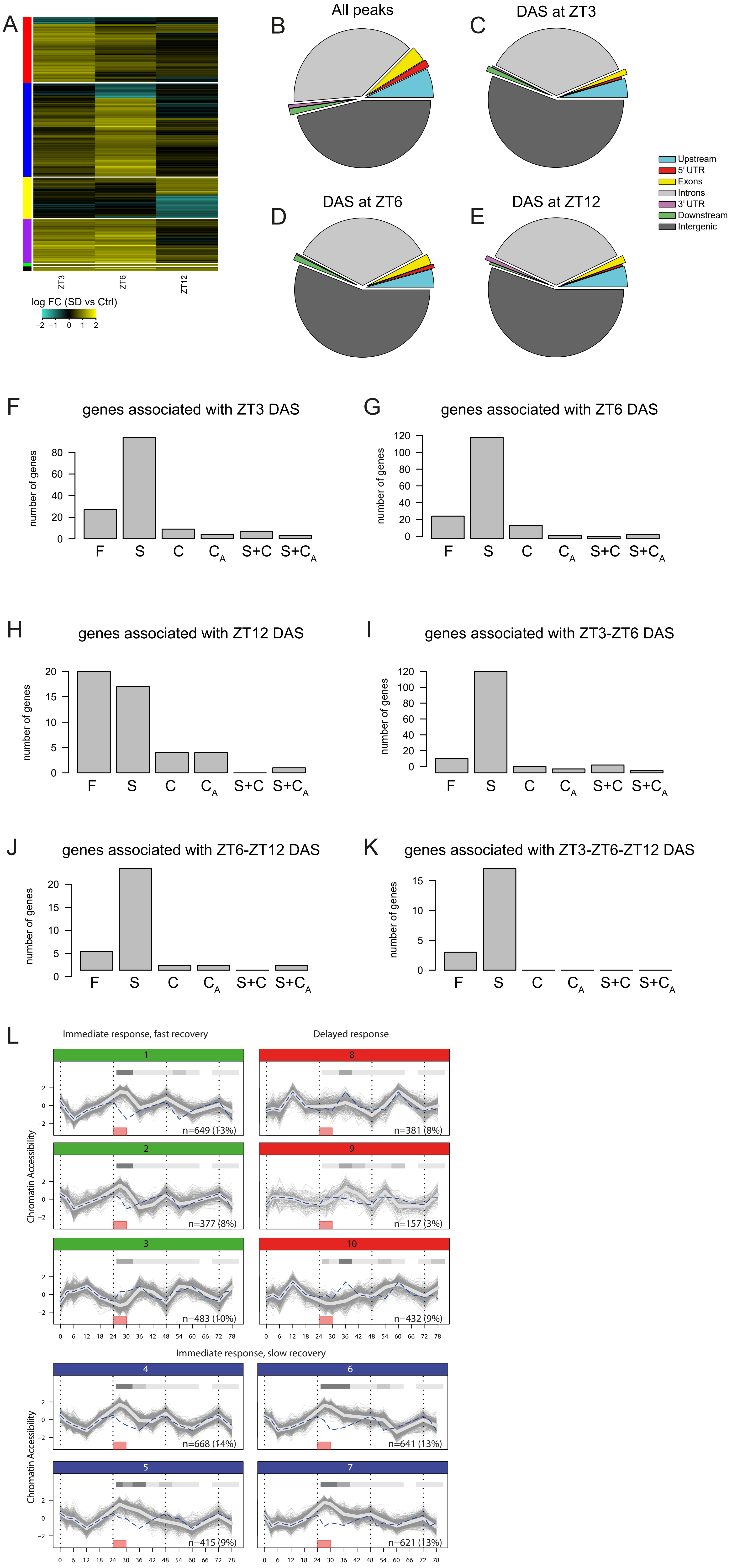


**Figure S3 ATAC-seq peak features. (A)** Heatmap of DAS for Ztime ZT3, ZT6 and ZT12. The heatmap was separated into 6 subgroups as follow: significant DAS uniquely detected between the time points T30 vs T3 (ZT3, red), T36 vs T6 (ZT6, blue), T42 vs T12 (ZT12, yellow). And significant DAS detected between multiple time points: T30 vs T3 AND T36 vs T6 (ZT3 & ZT6, purple), T36 vs T6 AND T42 vs T12 (ZT6 & ZT12, green), T30 vs T3 AND T36 vs T6 AND T42 vs T12 (ZT3 & ZT6 & ZT12, black). A hierarchical clustering was finally computed within each subgroups. **(B-E)** Genomic distribution of ATAC-seq peaks and DAS. **(F-K)** Repartition of genes associated with DAS among models at different times of day. **(L)** K-means clusters of the 3461 genes with statistically significant temporal gene expression (FDR adjusted p-value < 0.001, likelihood ratio test). Blue dashed line: average of the cluster under baseline, repeated for comparison over the three days of the experiment. Light grey thick line: cluster average. Red box: SD. Grey shaded bar at the top of each graph: proportion of genes with p-value <0.05 according to a likelihood ratio test between SD and baseline conditions at the same ZT.

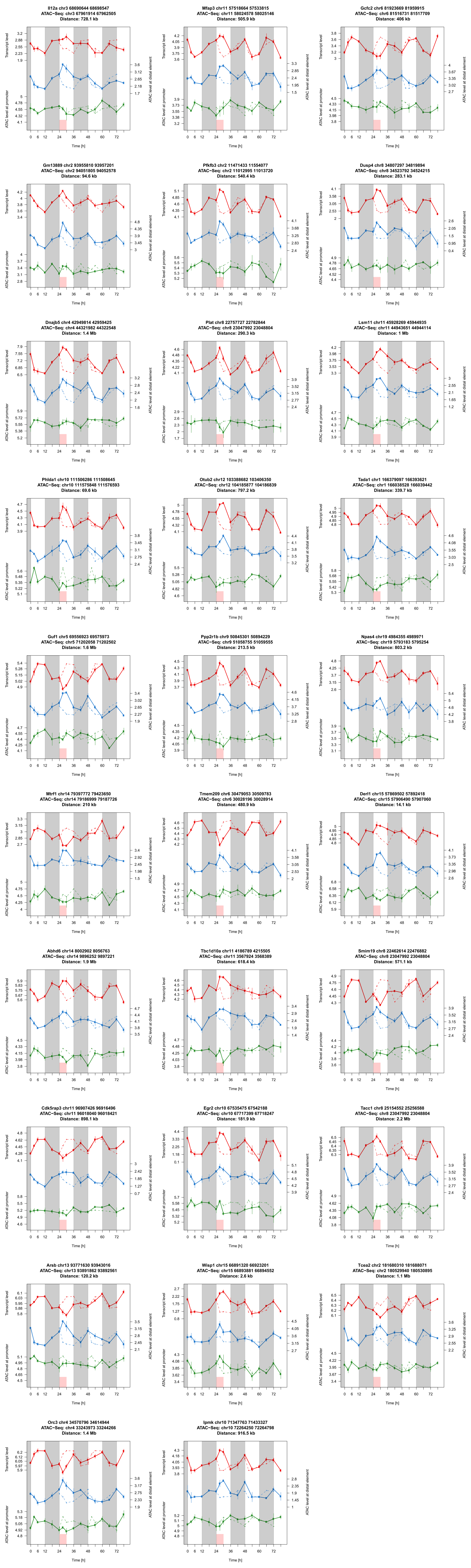


**Fig S4 Time courses of the distal elements, associated genes, and corresponding promoters**

Time courses of the distal elements, associated genes, and corresponding promoters involved in the top 34 associations (|ρ|>0.7), supplement to Fig 7. Header: gene name and genomic coordinates. ATAC-seq: coordinates of the top associated distal region. Distance in kb from the gene transcription start site to the closest boundary of the distal region. Red curve: gene expression level. Blue and green curves: accessibility level of distal region and promoter, respectively. Red box: SD.

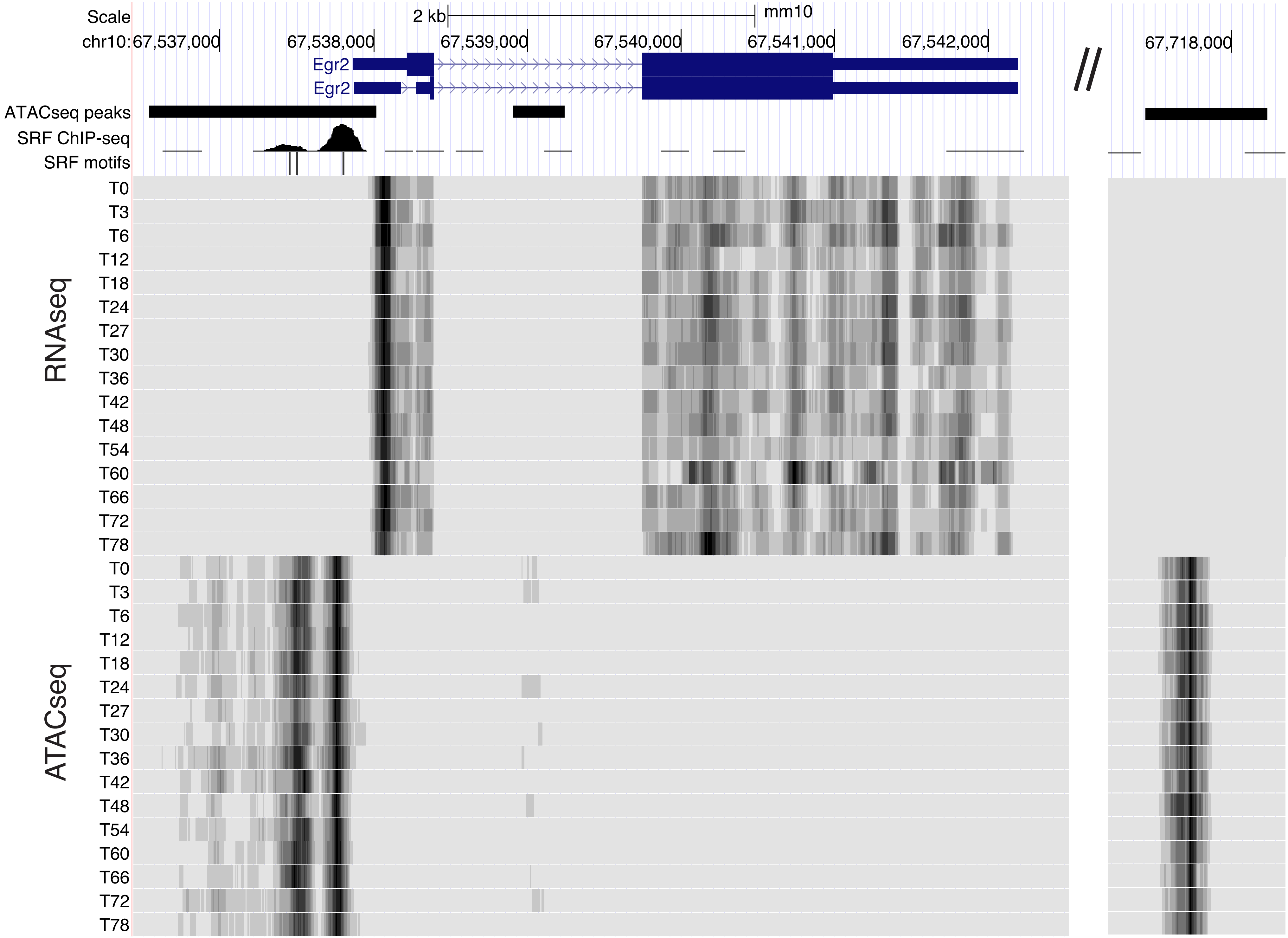


**Figure S5 Genome browser view** (genome.ucsc.edu) of RNAseq and ATACseq time course signal for Egr2, its promoter (left panel), and its interacting distal region (right panel). The "SRF ChIP-seq" track shows data from NIH3T3 cells, replicate 1 from Esnault Genes Dev 2014.

**Table S1.** **Transcript abundances for all detected genes used as input for model selection.**

Table S1 is provided as a separate dataset file.

**Table S2.** **Scaled transcript abundances used for k-means clustering.**

Table S2 is provided as a separate dataset file.

**Table S3 Tissue expression enrichment of genes overlapping with ATAC-seq positive regions**

| organId | organName | annotated | significant | expected | foldEnrichme | pValue | FDR |
|---|---|---|---|---|---|---|---|
| UBERON:0002313 | hippocampus pyramidal layer | 81 | 81 | 55.11 | 1.4697877 | 2.69E-14 | 1.71E-13 |
| UBERON:0014935 | cerebral cortex marginal layer | 219 | 218 | 149 | 1.46308725 | 1.70E-35 | 1.27E-34 |
| UBERON:0006083 | perirhinal cortex | 3532 | 3507 | 2403.03 | 1.4594075 | 0 | 0 |
| UBERON:0002191 | subiculum | 8630 | 8566 | 5871.51 | 1.45890921 | 0 | 0 |
| UBERON:0002728 | entorhinal cortex | 3577 | 3550 | 2433.65 | 1.45871428 | 0 | 0 |
| UBERON:0010415 | barrel cortex | 5813 | 5768 | 3954.93 | 1.45843289 | 0 | 0 |
| UBERON:0003883 | CA3 field of hippocampus | 3939 | 3905 | 2679.94 | 1.45712217 | 0 | 0 |
| UBERON:0013531 | retrosplenial region | 8956 | 8878 | 6093.31 | 1.45700777 | 0 | 0 |
| UBERON:0002739 | medial dorsal nucleus of thalamus | 8844 | 8763 | 6017.11 | 1.45634698 | 0 | 0 |
| CL:0000100 | motor neuron | 4064 | 4026 | 2764.98 | 1.4560684 | 0 | 0 |
| UBERON:0002667 | lateral septal nucleus | 9147 | 9059 | 6223.26 | 1.45566793 | 0 | 0 |
| UBERON:0002890 | anterior amygdaloid area | 9158 | 9069 | 6230.74 | 1.45552535 | 0 | 0 |
| UBERON:0001926 | lateral geniculate body | 8853 | 8766 | 6023.23 | 1.45536531 | 0 | 0 |
| UBERON:0004527 | alveolar process of maxilla | 903 | 894 | 614.37 | 1.45514918 | 2.78E-136 | 2.82E-135 |
| UBERON:0004725 | piriform cortex | 9214 | 9121 | 6268.84 | 1.45497413 | 0 | 0 |
| UBERON:0001875 | globus pallidus | 8888 | 8798 | 6047.04 | 1.45492671 | 0 | 0 |
| UBERON:0001883 | olfactory tubercle | 9069 | 8976 | 6170.19 | 1.4547364 | 0 | 0 |
| UBERON:0001989 | superior cervical ganglion | 8269 | 8184 | 5625.9 | 1.45470058 | 0 | 0 |
| UBERON:0005383 | caudate-putamen | 9413 | 9316 | 6404.23 | 1.45466356 | 0 | 0 |
| UBERON:0001935 | ventromedial nucleus of hypothalamus | 9091 | 8996 | 6185.16 | 1.45444904 | 0 | 0 |

**Table S4 BIC, weights of model fits and called model for all detected genes**

Table S4 is provided as a separate dataset file.

**Table S5 Associated genes and ATAC-seq peaks**

Table S5 is provided as a separate dataset file.
